# Myosin motor domains carrying mutations implicated in early or late onset Hypertrophic Cardiomyopathy have similar properties

**DOI:** 10.1101/622738

**Authors:** Carlos D. Vera, Chloe A. Johnson, Jonathan Walklate, Arjun Adhikari, Marina Svicevic, Srboljub M. Mijailovich, Ariana C. Combs, Stephen J. Langer, Kathleen M. Ruppel, James A. Spudich, Michael A. Geeves, Leslie A. Leinwand

**Affiliations:** BioFrontiers Institute and Department of Molecular, Cellular and Developmental Biology, University of Colorado Boulder, Boulder CO 80309 USA; School of Biosciences, University of Kent, Canterbury UK, CT2 7NJ; Faculty of Science, University of Kagujevac, Serbia; Department of Biology, Illinois Institute of Technology, Chicago IL 60616 USA; Stanford University School of Medicine, Department of Biochemistry, Stanford CA 94305 USA

**Keywords:** Human cardiomyopathies, hypertrophic cardiomyopathies, dilated cardiomyopathies, acto-myosin kinetics, kinetic modeling

## Abstract

Hypertrophic Cardiomyopathy (HCM) is a common genetic disorder that typically involves left ventricular hypertrophy and cardiac hypercontractility. Mutations in β cardiac myosin heavy chain (*β-MyHC*) are a major cause of HCM, but the specific mechanistic changes to myosin function that lead to the disease remain incompletely understood. Predicting the severity of any single *β-MyHC* mutation is hindered by a lack of detailed evaluation at the molecular level. In addition, since the cardiomyopathy can take 20 or more years to develop, the severity of the mutations must be somewhat subtle. We hypothesized that mutations which result in early onset disease may show more severe molecular changes in function compared to later onset mutations. In this work, we performed steady-state and transient kinetic analyses of myosins carrying 1 of 7 missense mutations in the motor domain. Of these 7, 4 have been identified in early onset cardiomyopathy screens. The derived parameters were used to model the ATP driven cross-bridge cycle. Contrary to our hypothesis, the results show no clear differences between early and late onset HCM mutations. Despite the lack of distinction between early and late onset HCM, the predicted occupancy of the force-holding actin.myosin.ADP complex at [Actin] = 3 *K*_*app*_ along with the closely related Duty Ratio (*DR*; fraction of myosin in strongly attached force-holding states) and the measured ATPases all change in parallel (in both sign and degree of change) compared to wild type (WT) values. Six of the 7 HCM mutations are clearly distinct from a set of DCM mutations previously characterized.

## Introduction

The most common inherited cardiovascular disease is HCM with a disease-prevalence of 1:250-500 (1, 2). Excluding those with a history of hypertension, aortic stenosis, other pre-existing systemic diseases, or being a world-class athlete, HCM is diagnosed as unexplained left ventricular hypertrophy and is typically accompanied by diastolic dysfunction (3). The first gene identified to contain a mutation leading to HCM was *MYH7* for the β-cardiac myosin heavy chain and was reported almost 30 years ago (4, 5). There are now thousands of mutations in genes that encode proteins of the cardiac sarcomere, and these account for 60-70% of HCM cases (6). There are 8 major sarcomeric proteins that are implicated in the majority of HCM index cases and their families, with 60-70% found in either β-cardiac myosin or myosin-binding-protein C (*MyBPC*). This suggests that myosin is an important target for therapeutic intervention (7, 8).

We consider here the difference between myosin mutations causing early onset versus late onset HCM. We hypothesized that β-cardiac myosin mutations associated with early onset HCM would be more severe than those mutations seen more typically in individuals who are diagnosed later in life. Analysis of the biochemical and biophysical properties of these 2 classes of myosins should reveal the severity of the mutational changes. We compared the properties of known adult pathogenic HCM mutations (R719W, R723G, and G741R) with novel sporadic mutations that had appeared in recent cardiomyopathy screens as specific to early onset patients (H251N, D382Y, P710R and V763M) (9, 10). We produced recombinant mutant and WT human myosin motors in differentiated C2C12 muscle cells, performed extensive kinetic analysis, and evaluated the severity of these mutations based on alterations to the cross-bridge cycle. We specifically focused on the cross bridge kinetics of the motor domain region utilizing the short sub-fragment 1 (sS1, corresponding to residues 1-808, Fig. S1) molecule.

The locations of the 7 residues in the β-MyHC protein under consideration here are shown in Fig. 1A. The high degree of conservation at those positions underscores the importance of these sites (Fig. 1B). Myosin is very vulnerable to mutations and there are now >400 different mutations described in *MYH7* (11, 12). A number of disease-causing *MYH7* mutations have been studied in the context of recombinant human *β-MyHC* motors (13–17). R403Q, R453C, R719W, R723G, G741R, and D239N are mutations that are widely recognized as pathogenic and have been seen in multiple families (18). Less prevalent sporadic mutations like H251N, D382Y, P710R, and V763M appeared in genetic screens of early-onset HCM patients (<15 years old) and their pathogenicity has not yet been clearly established (9, 18, 19).

**Figure 1.**
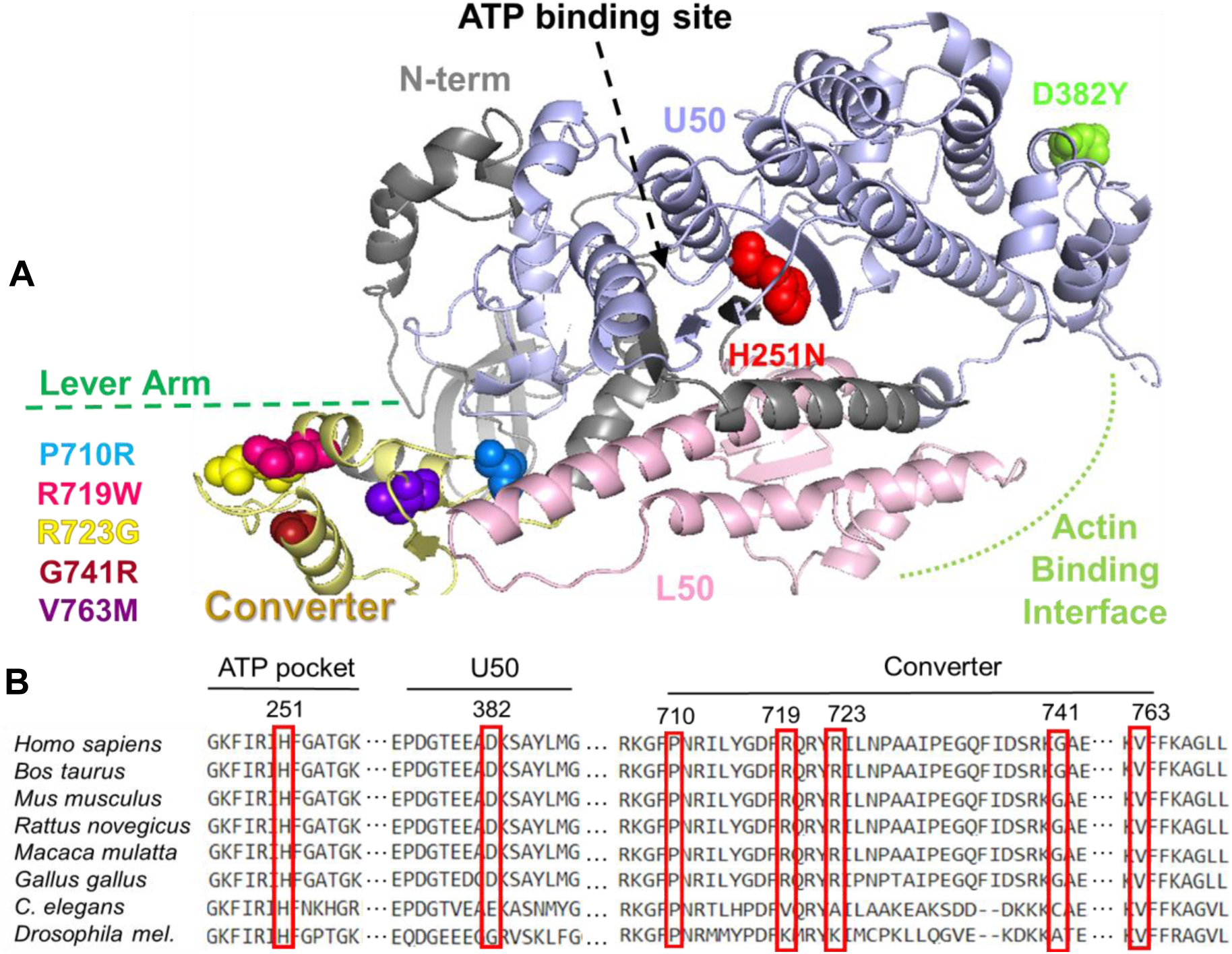
Structural location of HCM mutations. (A) Structural model of the catalytic domain of *β-MyHC* (based on PDB 4P7H). The heavy chain is shown as a ribbon diagram with the major subdomains and HCM mutations color-coded for the reader. The mutation sites are shown in space filling form in individual colors. The same colors are used throughout the figures and tables. Color code: Red -H251N; Green -D382Y; Light blue - P710R; Purple-V763M; Pink-R719W; Yellow-R723G; Maroon-G741R. Green dashed line represents the position of the lever arm, which was removed to clearly show the position of the mutated residues. (B) Alignment of conserved *MYH7* residues where mutations discussed throughout this study can be found.

H251N is in the central β-sheet (Fig.1A) that undergoes strain-induced twisting upon ATP-binding and communicates to the upper 50K domain to open and release actin. H251N was identified in a screen of 79 pre-adolescent children and later characterized biophysically (10, 17). Adhikari *et al.* found that this early onset mutation resulted in higher *k*_*cat*_, actin-gliding velocity, and intrinsic force (17). Located close to the cardiomyopathy loop of the actin binding domain, D382Y has been categorized as a variant of unknown (or uncertain) significance (VUS) (18, 19). Close to this residue is the well-studied pathogenic R403Q mutation which exhibits subtle biophysical changes compared to WT (14). P710, a residue on the border between the SH1 helix and the converter, has been reported to be mutated to an arginine, leucine or a histidine (10, 20). Unlike the arginine mutation, P710H and P710L have been seen multiple times, but the histidine is considered pathogenic, while the leucine and arginine mutations are rare and of unknown severity (10, 18, 20).

The myosin converter is well-known to be enriched for HCM-causing mutations with a range of adverse effects (21, 22). Converter mutations studied here include the late onset mutations R719W, R723G, and G741R and the early onset mutation V763M. Muscle biopsy and skinned fiber studies have shown the converter domain late onset mutations have increased fiber stiffness with subtle changes in cross-bridge kinetics (23–25). The steady-state *k*_*cat*_, the actin-gliding velocities, and the intrinsic force measurements of the myosin motor domain for these late onset pathogenic HCM mutations did not vary much when compared to WT (15). V763M has been reported in an early onset HCM screen and in a separate HCM cohort (19, 26). Both pathogenicity and age of symptom emergence are unclear for V763M (18).

We previously performed kinetic analysis of 5 late onset DCM mutations in *β-MyHC* (27). Overall, our analysis did not reveal a pattern of common defects in individual steps of the cycle, other than a few altered properties specific to the myosin subdomain where the mutation is localized. However, a pattern did emerge when the full cycle was modeled using the data from ATPase and stopped-flow experiments (28). Modeling predicted that the DCM mutations altered the steady-state motor function and the state occupancies in the minimal 8-step cross-bridge cycle in a manner consistent with a loss of the ability to generate steady-state force (27). Here, for the early- and late-onset HCM mutations, we show that the kinetic parameters of mutations also do not have a “unifying” disruption of the cycle. Contrary to our hypothesis, we did not detect any strong differences in kinetic parameters between mutations seen in late onset versus early onset patients, or among mutations in the different structural-domains of the motor, that account for the difference in the timing of pathological manifestation.

## Results

### ATPase activity

Steady-state ATPase data for 4 of the mutations examined here, R719W, R723G. G741R and V763M, have already been published (15, 17). These are listed in Table 1 together with the *k*_*cat*_ values for the remaining 4 mutations. The error on these measurements is of the order of 10% and 3 of the mutations, the early onset V763M and the late-onset R719W and R723G, differ from WT by less than 5%. The *k*_*cat*_ for G741R is less than 10% lower than WT. Three other mutations, H251N, D382Y and P710R, differ significantly from WT. The early onset H251N and D382Y are higher than WT by 24% and 20% respectively, while P710R is 39% lower than WT. Thus, there is no common pattern of change in the *k*_*cat*_ values for these HCM mutations and no difference in pattern between early and late onset groups. This lack of common patterns led us to focus on the transient kinetic analysis which can reveal more detail about how mutations change the ATPase cycle.

**Table 1.**
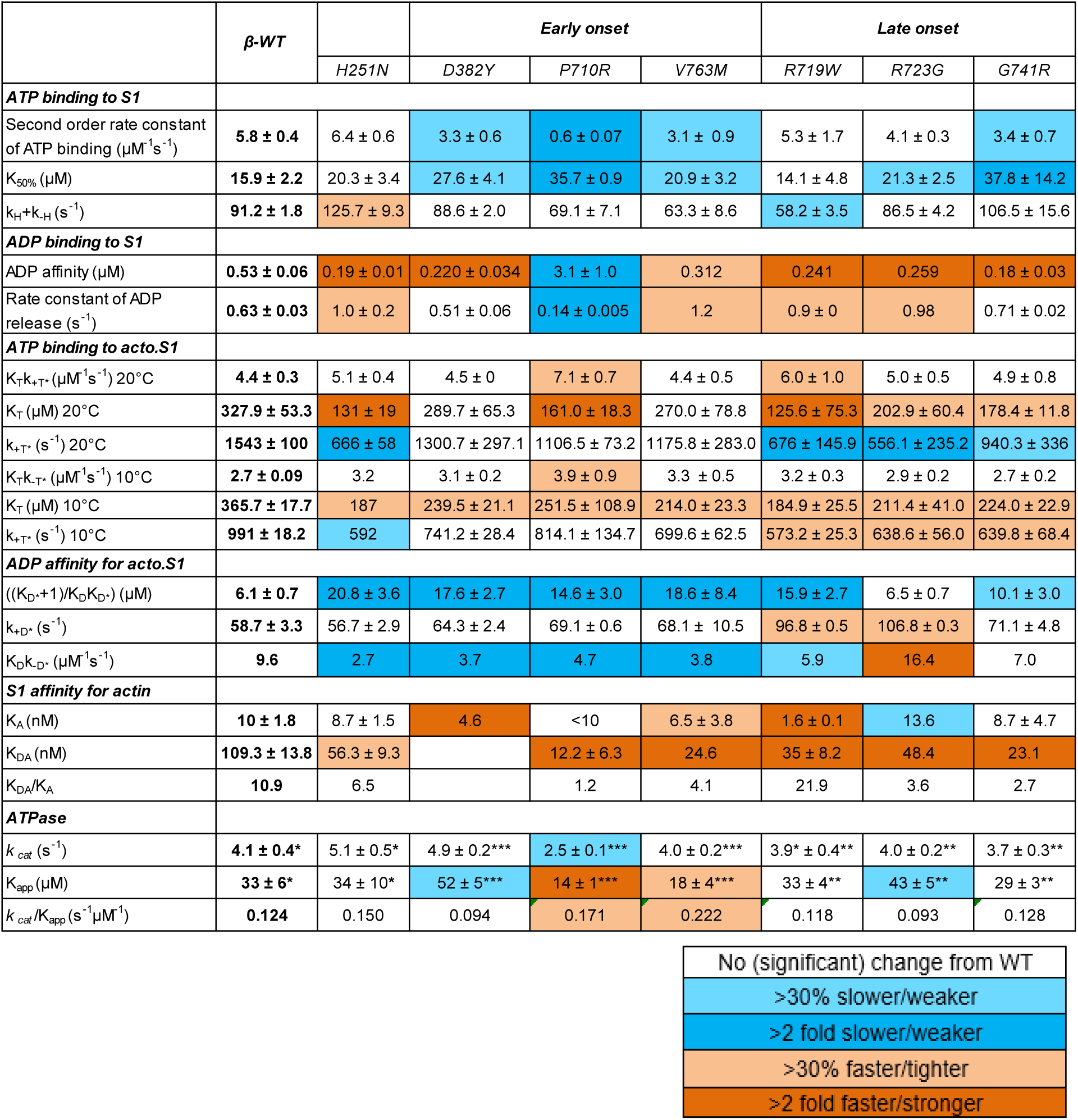
Kinetic parameters for J3-WT and 8 HCM mutations. The color code described in the smaller table to the right indicates degree of difference from WT. Experimental conditions for stopped-flow: 25 mM KCI, 5 mM MgCl2, 20 mM MOPS, pH 7.0, 20 C. Data are the mean ± S.E. values from 3-5 independent measurements with 4-6 technical replicates. Bottom of the table contains the ATPase values used for modeling: *ATPase values from (17). **ATPase values from (15). ***Unpublished ATPase values.

### Transient kinetics data summary

Considerable amounts of pre-steady state kinetic data have been collected for a number of DCM and HCM mutations in the *β-MyHC* motor domain. The descriptions of methods, analysis tools, model assumptions, data quality and details of the measurements have been presented in our earlier works (13, 14, 27–29). We have taken the same approach to understand the impact of this set of HCM mutations and to compare to the results of previous studies. Details of individual measurements and the evaluation of data quality are presented in the Supplementary Information (Figs. S2-S6). Here we focus on what effect each mutation has on the behavior of the cycle. For every mutant, each measurement was made at least 3 times with a minimum of 2 independent protein preparations. In general, all parameters were measured to a precision of better than 20% and in most cases better than 10%. Given this level of sensitivity, we assume any change of less than 20% is not significant.

The data are interpreted in terms of the 8-step ATP driven cross-bridge cycle that we have used previously (Fig. 2). Red shades indicate detached cross-bridges, yellow shades are weakly-attached, and blue shades represent strongly-attached force-holding cross-bridges. Table 1 shows the mean values and errors of the actin and nucleotide binding experiments for the steps in the cycle that are accessible, together with the ATPase parameters. To make the overall pattern of the induced changes clearer, the percent changes relative to WT are color coded in the table. The data are also summarized in Fig. 3, where the percentage differences in each parameter relative to WT are plotted.

**Figure 2.**
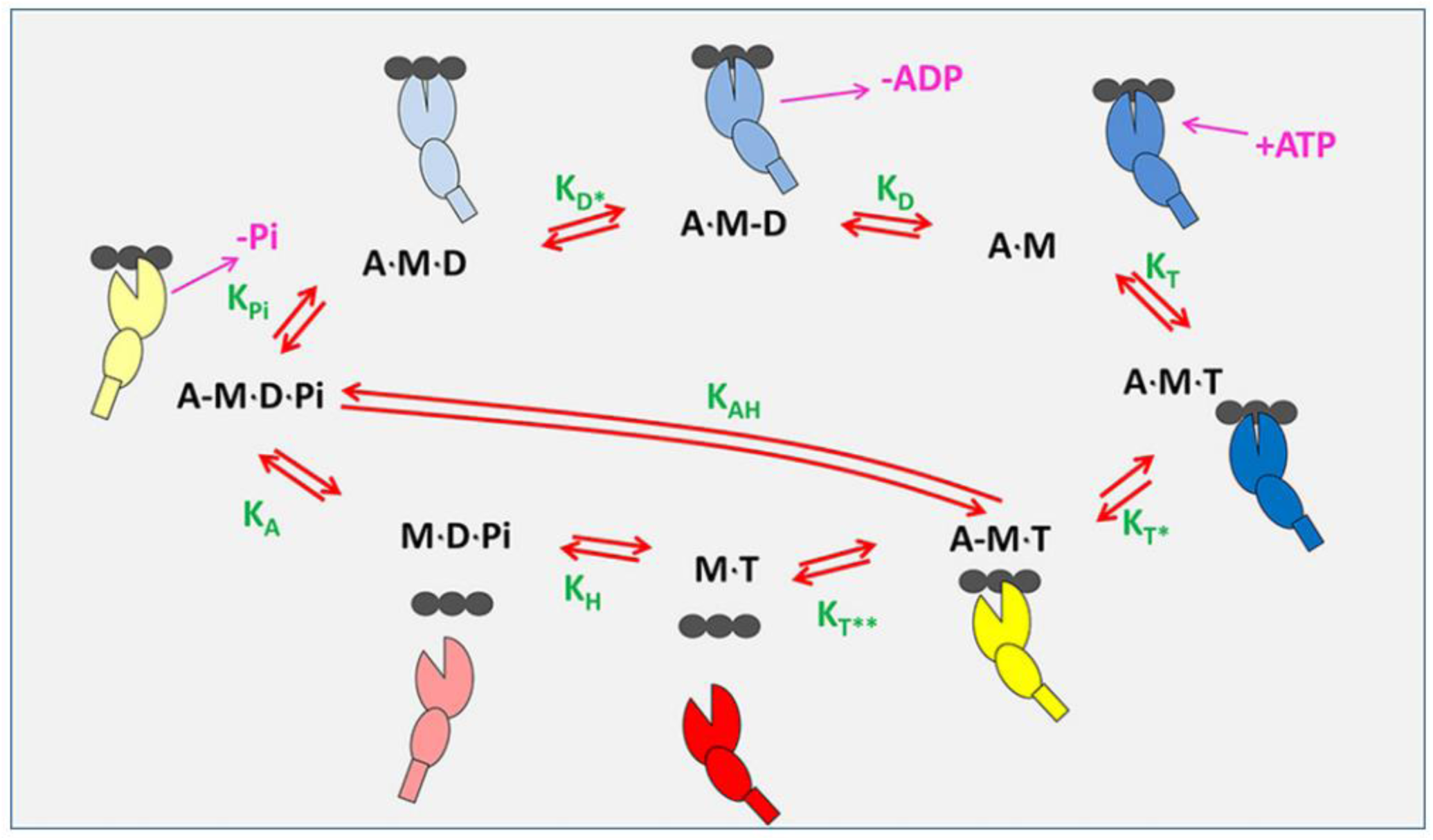
Actin⋅myosin ATPase-driven cross-bridge cycle. As discussed in Mijailovich et al, the basic ATPase cycle for myosin can be described in 8 steps: ATP binding, isomerization, ATP-induced dissociation, hydrolysis, Actin-attachment, phosphate-release + power stroke, 2^nd^ isomerization, and ADP release. The myosin is a composite of a large ellipse (motor domain), a smaller ellipse (converter), and a small square (lever arm) binding to an actin filament depicted as three black circles. A blue-shaded myosin is strongly-attached to actin (closed ellipse) and is progressively darker as it approaches the rigor state. The myosins with an open cleft away from actin are shown in yellow (weakly-attached) and red (detached). Nucleotide and P_i_ relevant steps are show in cerise, while green lettering indicates the equilibrium constant for the cycle step.

**Figure 3.**
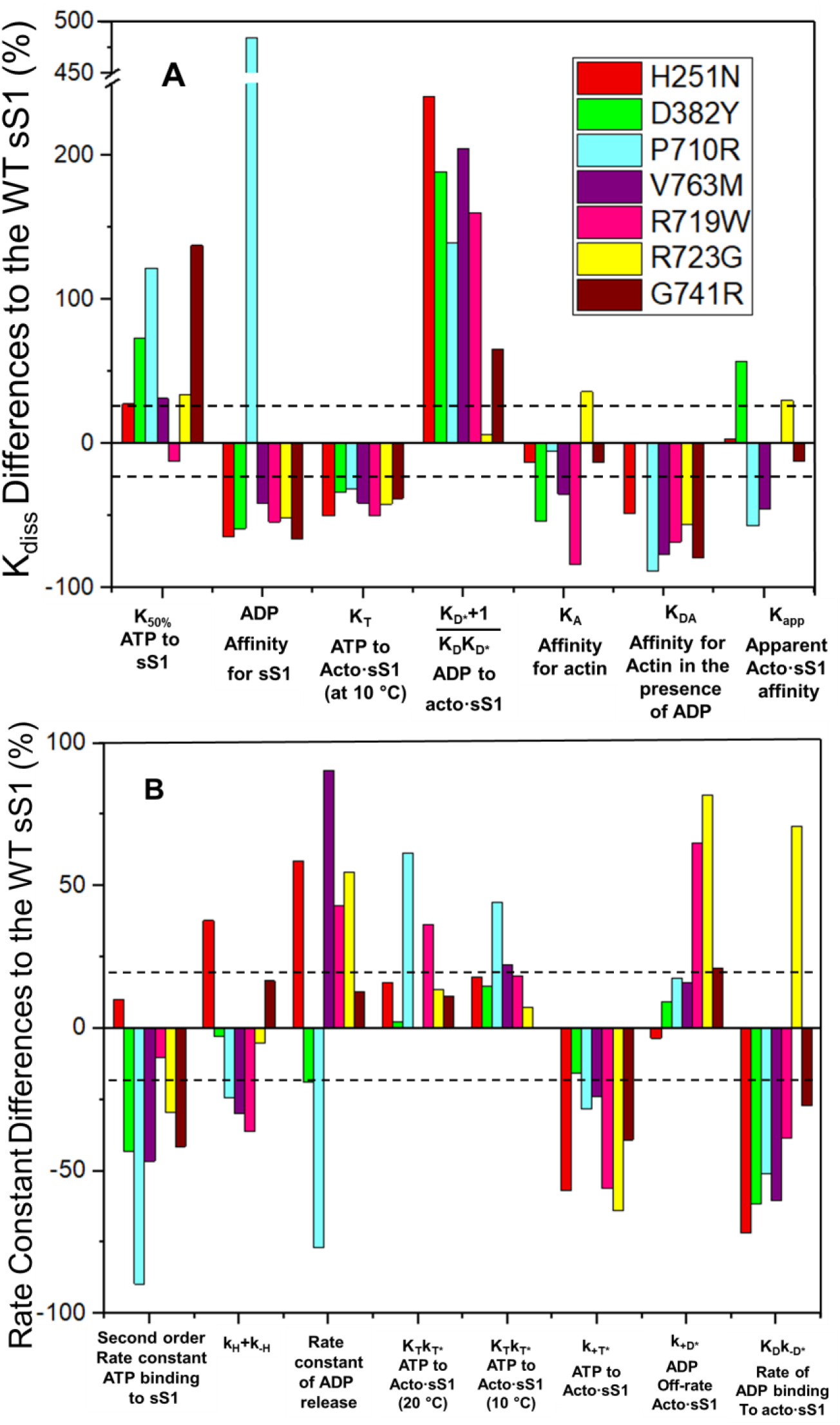
Summary of percentage differences in kinetic parameters of HCM mutations relative to WT. (A) %-change for measured dissociation or equilibrium constants. Also included is the apparent *K*_*app*_ from ATPase analysis. (B) %-change of several measured rate constants. Color-coded to match the parameter to each HCM mutation. The dashed lines represent ± 20% change in the parameter considered to be the precision of each measurement.

### Interaction of sS1 with nucleotide in the absence of actin

Even though the reactions are not part of the normal cycle, nucleotide binding to the motor in the absence of actin is measured to understand how the nucleotide binding pocket and the weakly-attached actin states might be affected by mutations. This process can be important to understand the cycle of the second myosin head, while the other can be attached to actin. The affinity of ATP for sS1 is weakened >2-fold for 2 mutations (P710R and G741R). All other mutations were 27-73% weaker except R719W which was indistinguishable from WT. In contrast, most mutants bound ADP >2-fold tighter, with the exception of V763M (just less than 2-fold tighter) and P710R (6-fold weaker). Thus, no simple pattern of behavior was apparent for the sS1 in the absence of actin, but most mutations significantly affected nucleotide binding.

### Interaction of sS1 with nucleotide in the presence of actin

The darker color coding of the data in Table 1 shows the parameters that change >2-fold (dark blue decrease, dark orange increase) and indicate that a large number of parameters have been affected, and at least one parameter for each mutant. Thus, the changes observed are not minor, despite relatively small changes in the value of the steady-state *k*_*cat*_.

Fig. 3A shows that in general, the value of *K*_*T*_ (the affinity of ATP for acto.sS1) measured at 10 °C where it is well defined, *K*_*A*_ (the affinity of actin for sS1), and *K*_*DA*_ (the affinity of actin for sS1.ADP) have tighter binding. For *K*_*T*_ this is >20% tighter, but none is as much as 2-fold tighter. For *K*_*DA*_ the affinities are mostly > 2 fold tighter with V763M, G741R and P710R > 4-fold tighter while the data for *K*_*A*_ is much more variable. The affinity of ADP for acto.sS1 is >2-fold weaker in 5 cases. R723G is an exception for both the *K*_*A*_ and ADP affinity for acto.sS1 parameters, but there is a general pattern.

Fig. 3B shows changes in measured rate constants. There is some consistent behavior, but it is not uniform for all mutations. The maximum rate constant for sS1 detachment from actin upon binding ATP (*k*_+*T*\*_) is generally slower (20-70%) and the rate constant for ADP binding to acto.sS1 is also generally slower (30-70%) with the exception of R723G which is increased by 70%. All other rate constants had variable changes or none at all. This highly variable pattern was also seen for the set of DCM mutations we previously reported (27).

### Modeling the ATPase cycle of HCM mutant motors

We modeled the complete cycle using all of our kinetic data following the approach outlined in recent papers comparing different myosin isoforms and different DCM-causing mutations in *MYH7* (27, 28). The transient kinetics data provide definition for 6 of the 8 steps of the cycle with a precision generally of ±20%. When combined with ATPase data (*k*_*cat*_ and *K*_*app*_), the full cycle allows us to predict acto⋅sS1 occupancy of states (Fig. 4; Supp. Table 3), along with key properties of the cycle: DR, shortening velocity, steady-state force (Fig. 5; Supp. Table 2), and ATP-economy (Fig. 7). Under isometric conditions in a muscle fiber, because of the mismatch of the thick and thin filament helicity, some myosin heads have readily accessible actin while others do not. We therefore modeled a range of actin concentrations ([A] = *K*_*app*_, *3 K*_*app*_, and *20 K*_*app*_, where the ATPase rate is 50, 75, and 98% of *k*_*cat*_ values, respectively) to facilitate comparison between the different mutant constructs under conditions that may match those of a contracting sarcomere. In addition, we modeled contraction under load as described previously (28), by assuming both ADP and P_i_ release steps were inhibited 3-fold under a 5-pN load for all mutations (see *Discussion*).

**Figure 4.**
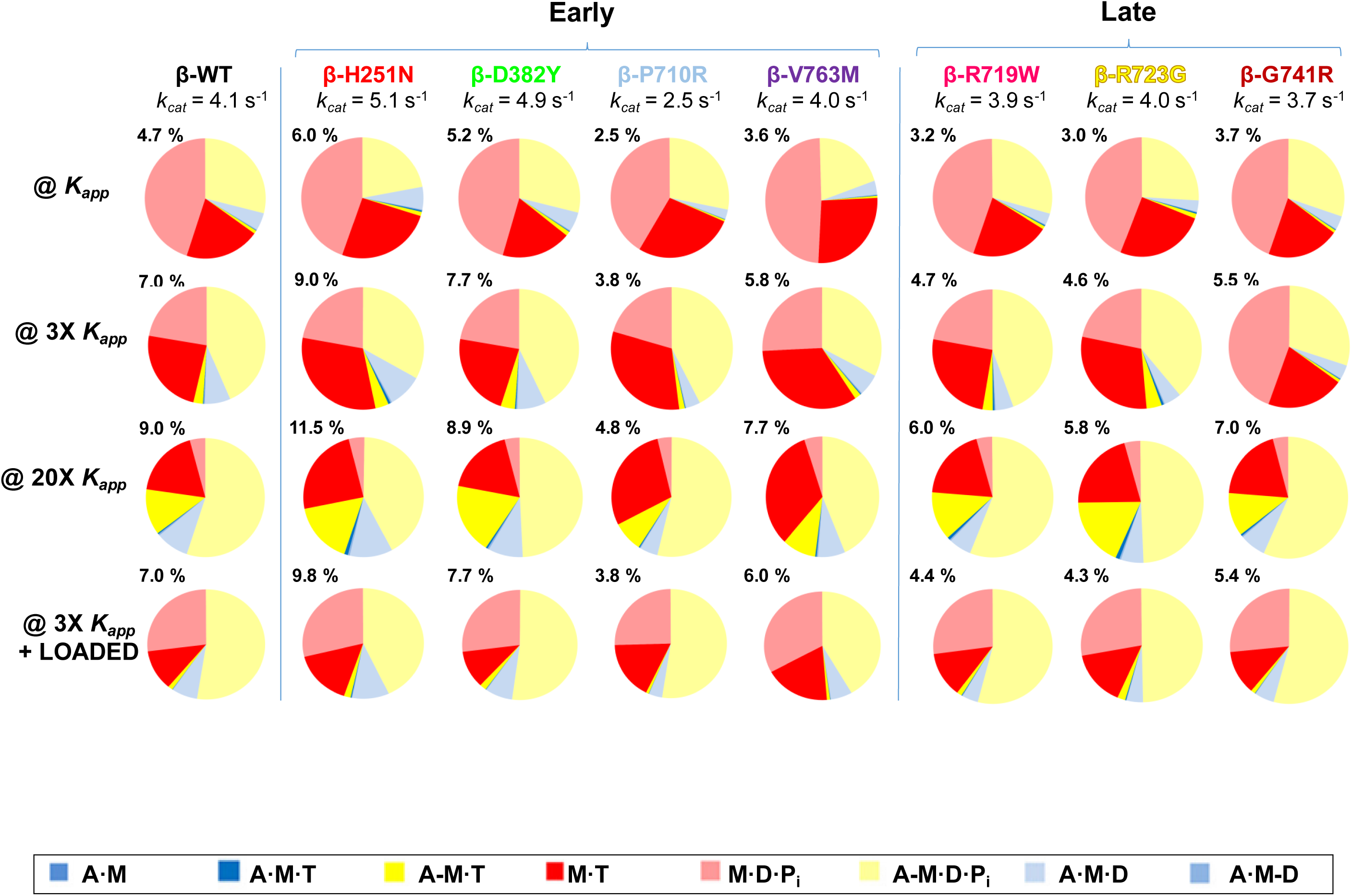
The occupancy of each intermediates from the 8-step ATPase cycle. The occupancy is shown at four actin-activated conditions; where [A] = *K*_*app*_, 3*K*_*app*_, 20*K*_*app*_ & 3*K*_*app*_ + 5 pN load. This figure is color-coded to match the scheme from Fig. 2 and the color legend shown below. Percentages at the upper left of each pie chart represents the percentage of time per cycle spent in the AMD state.

**Figure 5.**
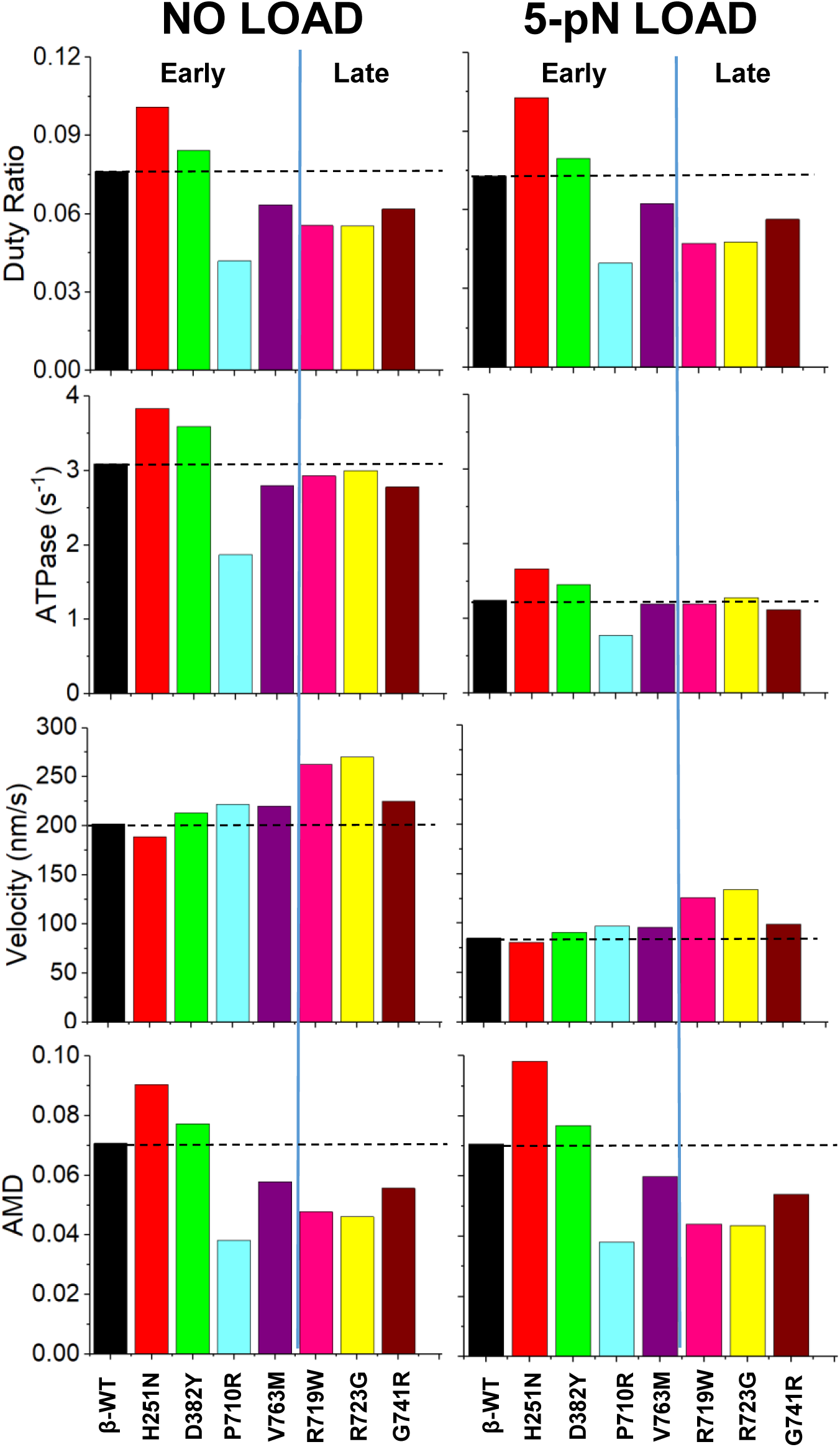
The calculated values for DR, *ATPase*, maximum shortening velocity, and occupancy of the force-holding A⋅M⋅D state for each of the HCM-causing mutations studied. The dashed line indicates the WT protein value. The left bar charts show the values based on kinetic measurements at [A] = 3*K*_*app*_. The right bar charts are the predicted results if the myosin additionally bears a 5-pN load.

The predicted occupancies are shown in Fig. 4. Note that the color schemes are the same in Fig. 2 and Fig. 4; red shades indicate detached cross-bridges, yellow shades are weakly-attached, and blue shades represent strongly-attached force-holding cross-bridges. The numbers above each pie chart represent the percentage of the force-holding A⋅M⋅D state (pale blue). The WT data predict at low actin concentration ([A] = *K*_*app*_), almost 75% of cross-bridges are detached with just 4.7% in the force-holding blue states, dominated by the A⋅M⋅D state. As actin concentration increases, the detached-states (red) decrease, the weakly-attached states (yellow) increase, and the strongly-attached A⋅M⋅D state increases to 9.0% at 20 *K*_*app*_. The application of a 5-pN load at *[A]* = *3 K*_*app*_ did not change the A⋅M⋅D state from 7.0% and increased the weakly-attached states (mainly A-M⋅D⋅P) at the expense of the detached M⋅T state. Note the occupancy values here are slightly lower than in (27) because of a slightly lower WT *k*_*cat*_ value used within this consistent data set.

Examining pie charts in Fig. 4 for the mutations shows that the general distribution of states is very similar between WT and H251N although the force-holding A⋅M⋅D state was higher (~25%) for H251N at all actin concentrations. This distribution was marginally larger (by about 10%) for D382Y. However, for P710R, V763M, R719W, R723G and G741R, the force-holding A⋅M⋅D states were smaller (by >20%) while the detached M⋅T states (deep red) were larger. These comparisons are easier to assess in Fig. 5 where the predicted A⋅M⋅D occupancy for [A] = 3 *K*_*app*_ with and without load are plotted along with the closely related *DR* and the calculated ATPases. All 3 parameters change in parallel (in both sign and degree of change) compared to the WT values.

The predicted velocity of shortening was estimated from the equation *V*_*o*_ = *d⋅ATPase/DR*, where *d* is the step size (assumed to be invariant at 5 nm). *V*_*o*_ shows little variation amongst the 7 mutants with 5 mutations varying by ~10% and only R719W and R723G increase velocity by >30% (Fig. 5). *In vitro* motility measurements have been published for several HCM mutations (15, 17). Unpublished data have been collected for the other HCM mutations studied here to consolidate with the kinetics data (Fig. 6, Supp. Table 8). Fig. 6 shows the mean velocity of the top 5% of smoothly moving filaments, normalized to the WT values, compared with our predictions. The normalization to the WT values allows comparison between different experimental conditions used. For 5 of the 7 mutations, the predicted velocities are in good agreement with those measured. There are large discrepancies for H251N (~50% higher than predicted) and P710R (>60% lower than predicted). We consider this discrepancy in the *Discussion*.

**Figure 6.**
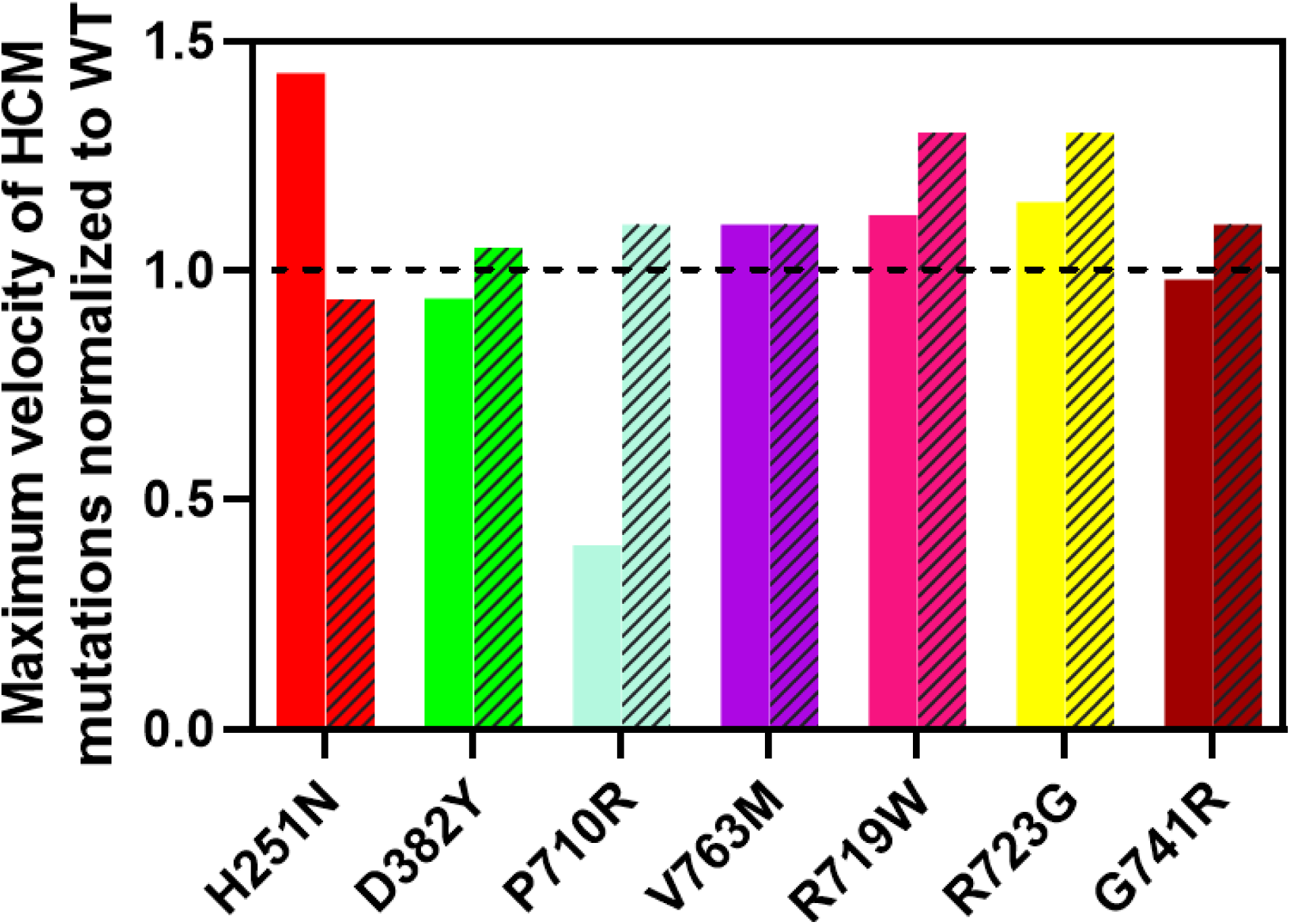
Comparison of normalized *in vitro* motility values to the predicted velocity of shortening. The solid coloring depicts the top 5% mean velocity values measured experimentally through unloaded *in vitro* motility. The bars with hatched patterns are the values predicted for the kinetic model analysis as described in (27). The dashed line indicates the normalized WT value of 1 for comparing the mutations.

The modeling data also allow an estimate of the economy of ATP usage, another parameter that could be involved in the development of HCM (30) if it results in a metabolic imbalance in the muscle. The predicted rates of ATP usage are shown in Fig. 7 for both a muscle fiber holding a 5-pN load (ATP used per pN) and when shortening at the maximum velocity with ATP being turned over at *k*_*cat*_ (ATP/nm moved). Once more, the changes in economy for each mutation, under both conditions reflected the changes in the ATPase rates relative to WT.

**Figure 7.**
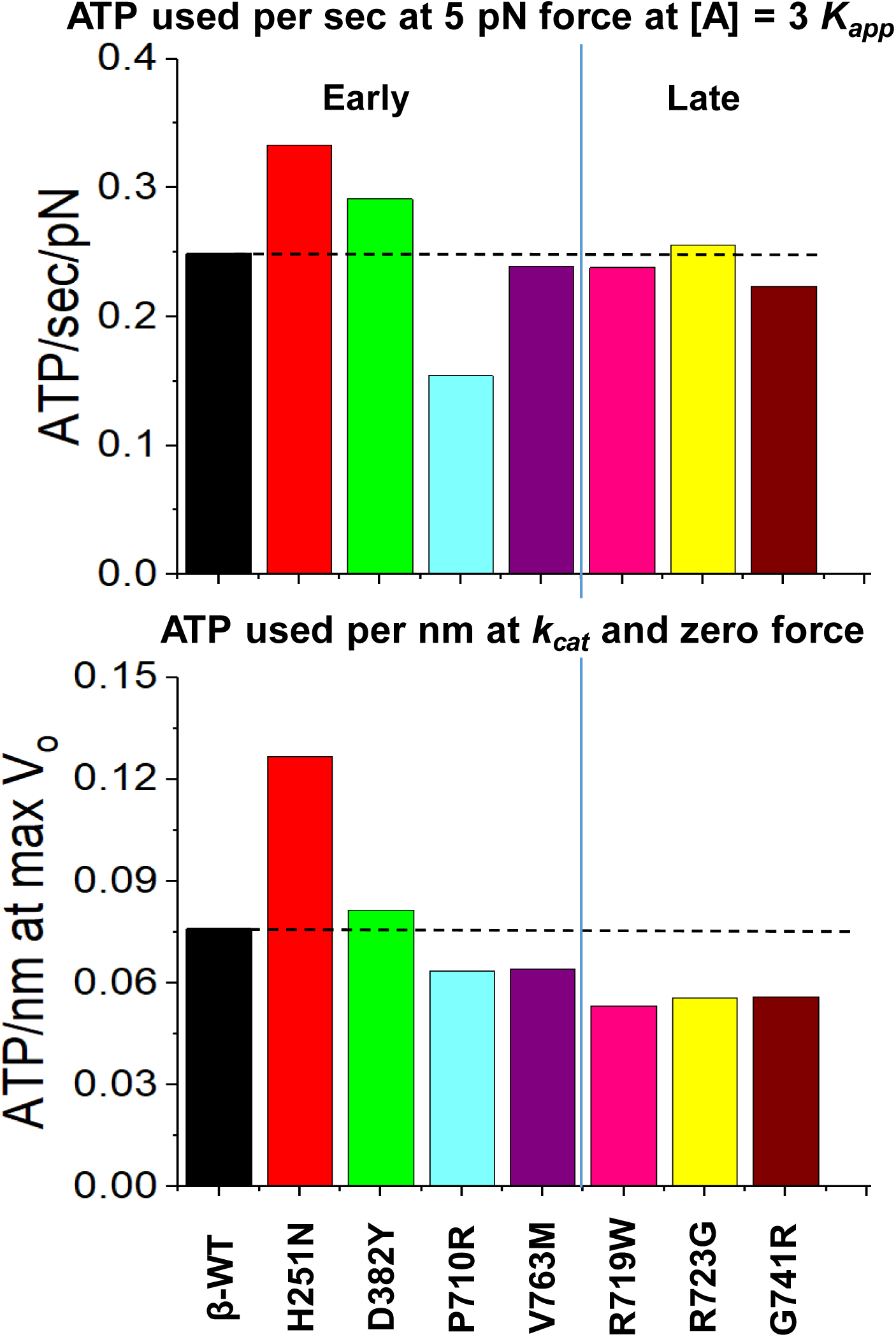
Economy of force generation and velocity for HCM mutations. Top, ATP usage per second per pN of force while generating 5 pN of force at [A] = 3*K*_*app*_. Bottom, ATP used per second at maximum shortening velocity (zero load). Because ATPase and *DR* have the same dependence on [A], this calculation is the same at all actin concentrations except very low values where the motor cannot maintain constant velocity.

## Discussion

We have presented here a detailed evaluation of the human β-cardiac myosin motor domain carrying 7 different HCM missense mutations. Before considering the implications of our data it is pertinent to point out the precision with which we can evaluate the data. All measured constants are defined to within an error of at least 20%, and in many cases better than 10%, using independent preparations of protein. The fitting of the data to the model uses all of the data to define the 5 missing constants in the 8-state cross-bridge cycle (Fig. 2). In our previous studies of DCM mutations (27) and the α and β cardiac isoforms (28), we showed the fitting was quite robust and that a ±20% change in any of the constants had minimal effect on the overall balance of the cycle except for the intermediate most directly affected by the varied constant. A similar analysis was carried out here (Supp. Table S5-6), so we are confident in the analysis of the cross-bridge cycles presented.

We compared our predictions of velocity with the measured velocities in in vitro motility assays using relative values of velocities (Fig. 6). Absolute values of velocity can be difficult to compare since these vary (see Table S8) between laboratories and within a single laboratory. Reference velocities are normally measured at the same time as novel measurements to allow comparisons. A similar problem can occur with *k*_*cat*_ measurements that depend critically on the quality of the protein preparation. Despite these difficulties our predictions are in good agreement with relative measured values for 5 of the 7 HCM mutations. The exceptions are H251N (which has a high *k*_*cat*_ and a high *DR* compared to WT) and P710R (which has low values of *k*_*cat*_ and *DR*). We noted previously for the DCM mutations that estimates of velocity can become problematic if the *DR* values are very low since higher myosin densities may be required to achieve the true maximum velocities.

Of the 7 mutations studied, 4 have been identified in children with HCM and 3 have mostly been reported in adults with HCM (Supp. Table S1). We previously reported a similar characterization of 2 additional late onset mutations, R403Q and R453C (13, 14, 27). The majority of mutations in myosin have been identified in adults and despite the mutant myosin being present in the heart during development and after birth, the cardiomyopathy can take 20-40 years to develop. We reasoned that early onset mutations may show more substantive changes in biophysical properties since they manifest earlier in life. The data presented here <u>*do not*</u> support this hypothesis. Each of the mutations (early onset and late onset) exhibited a set of changes in the measured parameters, at least 1 parameter by as much as ≥ 2-fold in each case. However, no general pattern of change was apparent that identified either the early onset group or the group as a whole. This remained true when the data were used to model the cross-bridge cycle (Figs. 4, 5, 7 Supp. Tables S2-4).

We previously identified common traits amongst myosins carrying 1 of 5 mutations associated with DCM (27). Most of the mutations were found to have a reduced *k*_*cat*_, lower occupancy of the force holding AMD state, a lower *DR*, and a more economical use of ATP for both rapid movement and force generation. Collectively, the DCM mutant myosins had an impaired capacity to generate or maintain force when working as an ensemble. Curiously, 1 of the mutations examined here shares these properties; the early onset P710R, suggesting some overlap between the groupings.

The remaining 6 mutations show mostly modest changes in properties as was previously observed for the R403Q and R453C HCM mutations. The *k*_*cat*_, economy of ATP usage and *DR’s* were similar to WT for D382Y, V763M, R719W, R723G and G741R while all 3 parameters were increased for H251N. Thus, for 6 of the 7 HCM mutations, the functional distinctions between HCM and DCM mutations that we previously reported remain valid. There are 3 perspectives that explain the molecular basis of HCM, including: (1) functional parameter changes; (2) increased number of available myosin heads; (3) and changes in load dependence of contractility (31). A combination of these subtle differences that are detected experimentally could help distinguish HCM severity among the different mutations.

There is considerable interest in the role of the interactions between the 2 motors of myosin in forming a down-regulated form of myosin: the Interacting Head Motif (IHM) (32, 33). Such motors are likely to be further stabilized by interactions with both the backbone of the thick filament and *MyBPC* (32–34). Myosin mutations that destabilize an off-state would make more heads available and lead to a hyper-contractile state, hypothesized to be the precursor to HCM (31–33, 35). The interaction between the off and on states of myosin is postulated to be regulated by phosphorylation of the *RLC* or *MyBPC*, mechanical strain on the thick filament, and possibly calcium (34, 36–39). Destabilization of the off-state could occur by reducing motor-motor, motor-backbone or motor-MyBPC interactions. Any effect on the interacting heads could override the modest effects we report here on the cross-bridge cycle for most of the HCM mutations. This seems less likely to be the case for P710R which appears similar to the DCM mutations where the large reduction in *k*_*cat*_ (30–40%) and *DR* (40-50%) will have a significant effect on the working cardiac muscle. Any effect on the interacting heads would need to be large enough to compensate for the loss of force-holding cross-bridges in the steady-state caused by these changes in the *DR*.

A recent study used the latest crystal structures docked into lower resolution EM images of the interacting motors to identify regions of the motor directly involved with the interaction. This work was combined with a molecular dynamics study to suggest how mutations might affect myosin function (40). HCM mutants were predicted to affect interacting heads, the stability of the M⋅D⋅P_i_ state required to form the interacting heads, the motor function, the stability of the motor, or some combination of the effects. From the Robert-Paganin *et al* study, our mutations H251N, R719W, R723G and G741R were predicted to affect motor function, which is consistent with our results, and to destabilize the sequestered state due to effects on IHM contacts, which is consistent with other experimental results (for a review, see (31)). P710R was predicted to alter motor function and destabilize the sequestered state by affecting the stability of the pre-power-stroke conformation, while V763M was predicted to affect mostly protein stability.

The question of what distinguishes mutations associated with early onset remains a valid question, but we need to address whether the diagnosis of early onset is a distinct group. It seems likely that factors such as genetic modifiers in addition to the myosin mutation contribute to many aspects of the disease, including age at which symptoms manifest (41, 42). In addition, the assays performed here are purely *in vitro* with single motor domains and other *in vitro* assays may shed further light on mechanisms of pathogenesis (31). While beyond the scope of this study, the assumption that the step size is unaffected could be tested using a single-molecule laser trap assay. Also, detailed direct biophysical measurements of the load dependence of the strongly-bound state are possible using Harmonic Force Spectroscopy (43) as well as measurements of the extent of sequestering of heads into a super-relaxed state (44). Furthermore, significant differences may be found using regulated thin filaments rather than purified actin in the ATPase measurements (14). In addition to these further *in vitro* studies, the mutant myosins may need a cellular or tissue environment to manifest their full pathological consequences.

At the outset of this series of studies, we began with the expectation that there would be common molecular pathways for HCM versus DCM mutations. The work has defined more sharply the ranges of effects mutations can have on the myosin motor and still have a functioning heart. To understand the way in which the mutations lead to myopathies, there is a need for more complex *in vitro* assays as well as assays that include cells and/or engineered 3D tissues to integrate the signals in the muscle which converge to cause disease. This includes the role of a heterogeneous mixture of WT and mutant myosins in tissue.

## Experimental Procedures

### Expression and purification of proteins

Producing recombinant β-cardiac myosin in C2C12 cells in several iterations, isoforms, and mutants has been described previously in (13, 14, 27, 29, 45, 46). The sS1 (residues 1-808) was followed by a flexible GSG (Gly-Ser-Gly) linker and either a C-terminal 8-residue (RGSIDTWV) PDZ-binding peptide or a C-terminal eGFP. The late onset mutations studied in (15) had an eGFP molecule at the C-terminus of the sS1 followed by the PDZ-binding peptide. The early-onset mutations lacked the eGFP molecule. Human ventricular essential light chain (MYL3 or ELC) with either an N-terminal 6X-His or a FLAG tag was co-expressed with the heavy chain. Over the course of the different experiments, a combination of affinity and ion exchange chromatography were used as in our previous studies (i.e. His or FLAG + Q column). The fast kinetics experiments in this study were done with protein purified with double affinity, a His-NTA resin and the PDZ-C-tag affinity system described in (47). WT-erbin PDZ was prepared as described in (47, 48) and crosslinked to the Sulfolink Coupling Resin (Thermo Fisher) using the manufacturer’s protocol. After the myosin was eluted from the His column, the sample was dialyzed into 1X TBS to remove the imidazole. The PDZ system uses 20-50 µM of “elution peptide” (NH_2_-WETWV-COOH from GenScript). After the second column, the myosin samples were buffer exchanged and frozen in the appropriate assay buffer. Actin preparations were made from bovine left ventricle with protocols adapted from (49, 50). The protocol for preparing pyrene-labelled actin is based on the methods described in (51).

### Steady-state actin-activated ATPase assays

All ATPase experiments were performed at 23°C room temperature in a buffer containing 10 mM imidazole, 3 mM MgCl_2_, 5 mM KCl, and 1 mM DTT at pH 7.5. A colorimetric assay was used to measure inorganic phosphate production at various time points and actin concentrations ranging 0 – 100 µM (52). The rates of phosphate production were plotted and fitted to the Michaelis-Menten equation using Origin (OriginPro) and/or GraphPad Prism to obtain the *k*_*cat*_ and *K*_*app*_.

### Transient kinetics

All stopped-flow measurements were performed at 20°C in 20 mM MOPS, 25 mM KCl, 5 mM MgCl_2_, 1 mM NaN_3_ at pH 7.0, unless indicated otherwise. Rapid-mixing experiments using 2-5 biological replicates, with 4-6 technical replicates, over a wide range of experimental substrates were performed in a High-Tec Scientific SF-61 DX2 stopped-flow system. Transient kinetic traces were initially fitted with TgK Scientific (Kinetic Studios) and subsequently plotted with Origin (OriginPro). For experiments probing the actin-myosin interaction with ATP and ADP we utilized the fluorescence signal for the pyrene-labelled actin, which has an excitation wavelength at 365-nm and the emission was detected after passing through KV389-nm cut-off filter. In the absence of actin, we relied on the intrinsic tryptophan fluorescence post-nucleotide binding where a tryptophan on the relay helix can be excited at 295-nm and the emission is detected with WG-320 filter.

### Modeling analysis

The parameters estimated experimentally by the transient kinetics analysis can be utilized to model the cross-bridge cycle by having a good estimate of the *k*_*cat*_ and *K*_*app*_ values obtained from the ATPase experiments and using the in-house MUSICO software (28). We previously reported a kinetic modeling analysis of 5 DCM mutations using this approach (27). This can be employed to predict the transient occupancy of the states in the myosin ATPase cycle, across a wide range of actin concentrations assuming initially a state where [ADP] = [P_i_] = 0, and the system proceeds to steady-state activity (28). Because the different steps of the cycle have interdependence to each other it is important that the experimental data provide a uniquely resolved set of modeling parameters to properly fit the rate and equilibrium constant of the cycle (Fig 5.4) (53). Consistent with our previous modeling work, the fitted parameters are defined to a precision of ~20%. Other well-defined assumptions and estimates applied to this mechano-chemical cycle model for β-MyHC are the isoform-specific constraint effect and the effect of load on the motor (54, 55). For a muscle fiber under isometric tension an approximate 5-pN of load with a 3-fold reduction in the ADP release rate constant are suggested as good estimates. This reduction of the ADP-release rate has little effect on the ATP flux, because this step is significantly faster than the overall *k*_*cat*_. Additionally, the rate of entry into the force-generating state is expected to be inhibited by load via the Fenn effect (56). Therefore, a 3-fold inhibition of the ATPase flux is applied to the model by reducing the entry rate into the force-holding state (A⋅M⋅D) which is *k*_*Pi*_ in our model (57). This assumes that a load will impact the Pi and ADP release steps similarly.

### Analysis of errors in the modeling

We have set out in our previous work the approach to defining the errors in the modeling (27, 28). The parameters that go into the model are all assumed to be defined to a precision of ~20% based on the estimate of errors of the measurements (see Table 1). Fitting the model to the ATPase data then produces definition of the remaining unmeasured parameters (actin affinity to M.ADP.Pi, *K*_*A*_; ATP hydrolysis, *K*_*H*_; ATP hydrolysis while attached to actin, *K*_*AH*_; the rate constant of Pi release/force generating step, *k*_*Pi*_; and the rate of reverse ATP dissociation, *k*_-*T*_). Thediagonals of resolution matrices, Fig. S3, demonstrate that the values are defined relatively independently of each other with the exception of *k*_*-T*_ which is codependent of the other values.

As previously, to assess the quality of the fits we varied each of the values in the Table S3 by ± 20% and refitted the data to assess the effect on the overall fitting. These have only a small effect on the other fitted parameters usually by much less than 20%. This can be seen in the analysis of the WT data in our previous publication (28). The results for the mutations used here were similar. Here, as we did for a set DCM mutations (27) we additionally varied *k*_*cat*_, the rate constant of the step limiting ADP release (*k*_*D\**_) and the hydrolysis step (*k*_*H*_) and as an illustration the data is shown for P710R and R723G in Table S6. In general, the test shows the fitting to be quite robust; a change in a parameter of 20% alters only those events most directly affected and to a similar extent as the 20% change imposed.

### In vitro motility

All the experiments were performed at 23°C. Glass coverslips (VWR micro cover glass) were coated with a mixture of 0.2% nitrocellulose (Ernest Fullam Inc.) and 0.2% collodion (Electron Microscopy Sciences) dissolved in amyl acetate (Sigma) and air-dried before use. A permanent double-sided tape (Scotch) was used to construct 4 channels on each slide, and 4 different experiments were performed on the same slide. Partially inactivated myosin heads in sS1 preparations were removed by the ‘dead-heading’ process before performing the motility assay. The process of ‘dead-heading’ had the following steps: A ten-fold molar excess of F-actin was added to myosin in the presence of 2 mM ATP; the mixture was incubated for 15 min in an ice bucket; 50 mM MgCl_2_ was added to form F-actin Mg^2+^-paracrystals and incubated for 5 min; the paracrystals were sedimented at 350,000g for 15 min; supernatant was collected, and the sS1 concentration was measured using the Bradford reagent (Bio-Rad). Before any experiments, dead-headed sS1 was diluted in 10% ABBSA [assay buffer (AB; 25 mM imidazole, pH 7.5, 25 mM KCl, 4 mM MgCl_2_, 1 mM EGTA, and 1 mM DTT) with bovine serum albumin (BSA, 0.1 mg/mL) diluted in AB], unless otherwise stated.

For motility experiments using pure actin, reagents were sequentially flowed through the channels in the following order: 10 μL of 4 µM SNAP-PDZ18 diluted in AB and incubated for 3 min; 20 µL of ABBSA to block the surface from nonspecific attachments and incubated for 2 min; 10 μL of a mixture of 8-residue (RGSIDTWV)-tagged human β-cardiac sS1 (~0.05 to 0.1 mg/mL) and incubated for 3 min; 20 μL of AB to wash any unattached proteins; and finally, 10 μL of the GO solution [5 to 10 nM tetramethylrhodamine (TMR)-phalloidin (Invitrogen)-labeled bovine actin, 2 mM ATP (Calbiochem), an oxygen-scavenging system consisting of 0.2% glucose, glucose oxidase (0.11 mg/mL; Calbiochem), and catalase (0.018 mg/mL; Calbiochem)], and an ATP regeneration system consisting of 1 mM phosphocreatine (Calbiochem) and creatine phosphokinase (0.1 mg/mL; Calbiochem) in ABBSA].

## ACKNOWLEDGEMENTS

J.A.S. is a co-founder of Cytokinetics and MyoKardia. L.A.L. is a co-founder of MyoKardia. J.A.S., L.A.L., and K.M.R. are members of the scientific advisory board for MyoKardia. The authors are supported by the following funding sources: Children’s Cardiomyopathy Foundation (L.A.L), National Institutes of Health 2R01GM029090 (M.A.G., L.A.L., C.D.V), National Institutes of Health 3R01GM029090-31S1 (C.D.V.), National Institutes of Health 1F31GM111058-01A1 (C.D.V) and National Institutes of Health 2R01HL117138-05 (L.A.L., J.A.S., C.D.V.). M.A.G., S.M.M., M.S. & J.W. received funding from the European Union’s Horizon 2020 Research and Innovation Programme grant No. 777204 SILICOFCM. We also thank Sam Lynn for technical assistance

## AUTHOR CONTRIBUTIONS

M.A.G. and L.A.L. conceived the study and supervised each step of the work. C.D.V. designed, performed and analyzed steady-state and pre-steady state experiments, and was responsible for cell culture, adenovirus and protein preparations. C.A.J. and J.W. and designed and performed stopped-flow experiments as well as initiating most of the kinetic modeling analysis. C.A.J., S.M.M., and M.S. completed the kinetic modeling. A.C.C. and S.J.L. provided substantial technical support in cell culture. A.S.A., K.M.R., and J.A.S. designed and performed ATPase, *in vitro* motility assays and the respective data analysis.

