## Supplementary Information for "Myosin motor domains carrying mutations implicated in early or late onset Hypertrophic Cardiomyopathy have similar properties"

Running title: Early and late onset HCM causing mutations in MYH7

*To whom correspondence should be addressed: Michael A. Geeves School of Biosciences, University of Kent, Canterbury, UK, CT2 7NJ; Leslie A. Leinwand BioFrontiers Institute and/or Department of Molecular, Cellular and Developmental Biology, University of Colorado, Boulder CO 80309

**Table S1**. This table describes a set of queries regarding the clinical frequency of HCM linked or associated mutations in MYH7 assessed using the NCBI tool ClinVar and the Variant Effect Predictor (VEP) tool for Ensembl (1, 2). H251N, P710R, P710L and V763M are predicted to have physico-chemical effects and possibly be disease-causing because those residues are highly conserved, but due to rarity it is not enough to be considered pathogenic, hence being categorized “Deleterious, VUS”.

| MYH7 Mutation | ClinVar Variation ID | Has the mutation been described in more than one person? | Has the mutation been seen in multiple families? | Has the mutation affected associated family members of the patient? | Is it predicted to be pathogenic? PolyPhen - A SIFT – B Condel - C | Is it present in large population databases? | Identified as early onset? (<15 y.o.) |
|---|---|---|---|---|---|---|---|
| H251N | n/a | No | No | No | Deleterious, VUS | No | Yes |
| D382Y | 237432 | Yes | No | No | A: ✓ B: ✓ C: ✓ | No | Yes |
| R403Q | 14087 | Yes | Yes | Yes | A: ✓ B: ✓ C: ✓ | No | No |
| R453C | 14089 | Yes | Yes | Yes | A: ✓ B: ✓ C: ✓ | No | No |
| P710R | n/a | No | No | No | Deleterious, VUS | No | Yes |
| P710H* | 177734 | Yes | No | No | A: ✓ B: ✓ C: ✓ | No | No |
| P710L* | 177673 | Yes | No | No | Deleterious, VUS | No | No |
| R719W | 14104 | Yes | Yes | Yes | A: ✓ B: ✓ C: ✓ | No | No |
| R723G | 42885 | Yes | Yes | Yes | A: ✓ B: ✓ C: ✓ | No | No |
| G741R | 42890 | Yes | Yes | Yes | A: ✓ B: ✓ C: ✓ | No | No |
| V763M | 177642 | Yes | No | n.d. | Deleterious, VUS | No | Yes |

* = Reported mutations on residue P710, however these were not characterized in this work.

✓ = Algorithm predicts pathogenic and/or “likely-pathogenic”.

**Table S2. Fitted Rate and Equilibrium Constants of the ATPase cycle.** Highlighted to show which fitted & which measured and which derived from assumption or detailed balance.

A. Equilibrium Constants

| | *Units* | *β-WT* | *H251N* | *D382Y* | *P710R* | *V763M* | *R719W* | *R723G* | *G741R* |
|---|---|---|---|---|---|---|---|---|---|
| $K_{app}$ | μM | 33 | 34 | 52 | 14 | 18 | 33 | 43 | 29 |
| $k_{cat}$ | $s^{-1}$ | 4.1 | 5.1 | 4.9 | 2.5 | 4 | 3.9 | 4 | 3.7 |
| $K_A$ | $\mu M^{-1}$ | 0.0199 | 0.0148 | 0.0124 | 0.0501 | 0.0306 | 0.0204 | 0.014 | 0.0235 |
| $K_{pi}$ | mM | 100 | 100 | 100 | 100 | 100 | 100 | 100 | 100 |
| $K_{D*}$ | | 0.167 | 0.167 | 0.167 | 0.167 | 0.167 | 0.167 | 0.167 | 0.167 |
| $K_D$ | μM | 36 | 36 | 36 | 36 | 36 | 36 | 36 | 36 |
| $K_T$ | $\mu M^{-1}$ | 0.0036 | 0.00763 | 0.00345 | 0.0062 | 0.0037 | 0.00796 | 0.0049 | 0.0056 |
| $K_{T*}$ | | 154 | 66.6 | 130 | 110.65 | 117.58 | 67.6 | 55.6 | 94 |
| $K_{T**}$ | μM | 1000 | 1000 | 1000 | 1000 | 1000 | 1000 | 1000 | 1000 |
| $K_H$ | | 9.4 | 8.6 | 11.7 | 4.9 | 6.5 | 8.5 | 7.2 | 8.4 |
| $K_{AH}$ | | 66.6 | 61.8 | 35.5 | 34.7 | 46.5 | 60.7 | 51.4 | 60.1 |

B. Forward Rate Constants

| | Units | β-WT | H251N | D382Y | P710R | V763M | R719W | R723G | G741R |
|---|---|---|---|---|---|---|---|---|---|
| $k_A$ | $\mu M^{-1}s^{-1}$ | 19.9 | 14.8 | 12.4 | 50.1 | 30.6 | 20.4 | 14 | 23.5 |
| $k_{Pi}$ | $s^{-1}$ | 7.1 | 11.6 | 8.4 | 4.4 | 8.6 | 6.6 | 7.7 | 6.2 |
| $k_{D*}$ | $s^{-1}$ | 59 | 56.7 | 64.3 | 69 | 68 | 96.8 | 106.8 | 71.1 |
| $k_D$ | $s^{-1}$ | 1000 | 1000 | 1000 | 1000 | 1000 | 1000 | 1000 | 1000 |
| $k_T$ | $\mu M^{-1}s^{-1}$ | 10.17 | 10.07 | 10.32 | 10.25 | 10.15 | 10.03 | 10.07 | 10.02 |
| $k_{T*}$ | $s^{-1}$ | 1540 | 666 | 1300 | 1106.5 | 1175.8 | 676 | 556 | 940 |
| $k_{T**}$ | $s^{-1}$ | 1000 | 1000 | 1000 | 1000 | 1000 | 1000 | 1000 | 1000 |
| $k_H$ | $s^{-1}$ | 13.1 | 12.1 | 16.4 | 6.8 | 9.1 | 11.9 | 10.0 | 11.8 |
| $k_{AH}$ | $s^{-1}$ | 13.3 | 12.35 | 7.1 | 6.9 | 9.3 | 12.1 | 10.3 | 12.0 |

C. Backward Rate Constants

| | ***Units*** | ***β-WT*** | ***H251N*** | ***D382Y*** | ***P710R*** | ***V763M*** | ***R719W*** | ***R723G*** | ***G741R*** |
|---|---|---|---|---|---|---|---|---|---|
| $k_{-A}$ | $s^{-1}$ | 1000 | 1000 | 1000 | 1000 | 1000 | 1000 | 1000 | 1000 |
| $k_{-Pi}$ | $mM^{-1}s^{-1}$ | 0.071 | 0.116 | 0.084 | 0.044 | 0.086 | 0.066 | 0.077 | 0.062 |
| $k_{-D*}$ | $s^{-1}$ | 353.29 | 339.52 | 385.03 | 413.17 | 407.19 | 579.64 | 639.52 | 425.75 |
| $k_{-D}$ | $\mu M^{-1}s^{-1}$ | 27.78 | 27.78 | 27.78 | 27.78 | 27.78 | 27.78 | 27.78 | 27.78 |
| $k_{-T}$ | $s^{-1}$ | 2824.1 | 1319.4 | 2989.9 | 1653.5 | 2743.5 | 1260.2 | 2054.3 | 1789.8 |
| $k_{-T*}$ | $s^{-1}$ | 10 | 10 | 10 | 10 | 10 | 10 | 10 | 10 |
| $k_{-T**}$ | $\mu M^{-1}s^{-1}$ | 1 | 1 | 1 | 1 | 1 | 1 | 1 | 1 |
| $k_{-H}$ | $s^{-1}$ | 1.4 | 1.4 | 1.4 | 1.4 | 1.4 | 1.4 | 1.4 | 1.4 |
| $k_{-AH}$ | $s^{-1}$ | 0.2 | 0.2 | 0.2 | 0.2 | 0.2 | 0.2 | 0.2 | 0.2 |

**Table S3. Predicted cross-bridge cycle parameters at three actin concentrations from the modelled ATPase cycle.**

| *Isoform* | *[Actin] (μM)* | *ATPase ($s^{-1}$)* | *Detached* | *Weakly attached* | *Strongly attached* | *Duty Ratio* | *Velocity (μm/s)* |
|---|---|---|---|---|---|---|---|
| ***β-WT*** | | | | | | | |
| | $K_{app}$ = 33 | 2.06 | 0.65 | 0.30 | 0.05 | 0.05 | 0.20 |
| | 3 $K_{app}$ = 99 | 3.09 | 0.46 | 0.46 | 0.08 | 0.08 | 0.20 |
| | 20 $K_{app}$ = 660 | 3.91 | 0.23 | 0.68 | 0.10 | 0.10 | 0.20 |
| | 3 $K_{app}$ + Load | 1.25 | 0.39 | 0.54 | 0.07 | 0.07 | 0.09 |
| **H251N** | | | | | | | |
| | $K_{app}$ = 34 | 2.56 | 0.70 | 0.23 | 0.07 | 0.07 | 0.19 |
| | 3 $K_{app}$ = 102 | 3.83 | 0.53 | 0.37 | 0.10 | 0.10 | 0.19 |
| | 20 $K_{app}$= 680 | 4.87 | 0.28 | 0.59 | 0.13 | 0.13 | 0.19 |
| | 3 $K_{app}$ + Load | 1.67 | 0.45 | 0.45 | 0.10 | 0.10 | 0.08 |
| **D382Y** | | | | | | | |
| | $K_{app}$ = 52 | 2.43 | 0.64 | 0.30 | 0.06 | 0.06 | 0.21 |
| | 3 $K_{app}$ = 156 | 3.59 | 0.45 | 0.47 | 0.08 | 0.08 | 0.21 |
| | 20 $K_{app}$ = 1040 | 4.14 | 0.22 | 0.68 | 0.10 | 0.10 | 0.21 |
| | 3 $K_{app}$ + Load | 1.46 | 0.38 | 0.54 | 0.08 | 0.08 | 0.09 |
| **P710R** | | | | | | | |
| | $K_{app}$ = 14 | 1.24 | 0.68 | 0.29 | 0.03 | 0.03 | 0.22 |
| | 3 $K_{app}$ = 42 | 1.87 | 0.52 | 0.44 | 0.04 | 0.04 | 0.22 |
| | 20 $K_{app}$ = 280 | 2.36 | 0.33 | 0.62 | 0.05 | 0.05 | 0.22 |
| | 3 $K_{app}$ + Load | 0.77 | 0.43 | 0.53 | 0.04 | 0.04 | 0.10 |

| | | | | | | | |
|---|---|---|---|---|---|---|---|
| **V763M** | | | | | | | |
| | $K_{app}$ = 18 | 1.75 | 0.75 | 0.21 | 0.04 | 0.04 | 0.22 |
| | 3 $K_{app}$ = 54 | 2.80 | 0.59 | 0.34 | 0.06 | 0.06 | 0.22 |
| | 20 $K_{app}$ = 360 | 3.75 | 0.38 | 0.53 | 0.09 | 0.09 | 0.22 |
| | 3 $K_{app}$ + Load | 1.20 | 0.51 | 0.43 | 0.06 | 0.06 | 0.10 |
| **R719W** | | | | | | | |
| | $K_{app}$ = 33 | 1.95 | 0.66 | 0.30 | 0.04 | 0.04 | 0.26 |
| | 3 $K_{app}$ = 99 | 2.93 | 0.47 | 0.47 | 0.06 | 0.06 | 0.26 |
| | 20 $K_{app}$ = 660 | 3.71 | 0.24 | 0.69 | 0.07 | 0.07 | 0.26 |
| | 3 $K_{app}$ + Load | 1.19 | 0.56 | 0.40 | 0.05 | 0.05 | 0.13 |
| **R723G** | | | | | | | |
| | $K_{app}$ = 43 | 2.00 | 0.69 | 0.27 | 0.04 | 0.04 | 0.27 |
| | 3 $K_{app}$ = 129 | 3.00 | 0.51 | 0.43 | 0.06 | 0.06 | 0.27 |
| | 20 $K_{app}$ = 860 | 3.81 | 0.25 | 0.68 | 0.07 | 0.07 | 0.26 |
| | 3 $K_{app}$ + Load | 1.28 | 0.43 | 0.52 | 0.05 | 0.05 | 0.13 |
| **G741R** | | | | | | | |
| | $K_{app}$ = 29 | 1.85 | 0.65 | 0.31 | 0.04 | 0.04 | 0.23 |
| | 3 $K_{app}$ = 87 | 2.78 | 0.47 | 0.47 | 0.06 | 0.06 | 0.22 |
| | 20 $K_{app}$ = 580 | 3.52 | 0.24 | 0.68 | 0.08 | 0.08 | 0.22 |
| | 3 $K_{app}$ + Load | 1.12 | 0.39 | 0.55 | 0.06 | 0.06 | 0.10 |

**Table S4. Predicted Occupancy of the 8 states during the ATPase cycle.**

| *Isoform* | *[Actin] (μM)* | *A·M* | *A·M·T* | *A-M·T* | *M·T* | *M·D·Pi* | *A-M·D·Pi* | *A·MD* | *A·M-D* |
|---|---|---|---|---|---|---|---|---|---|
| ***β-WT*** | | | | | | | | | |
| | $K_{app}$ = 33 | 0.00012 | 0.0014 | 0.0086 | 0.20 | 0.45 | 0.29 | 0.047 | 0.0021 |
| | 3 $K_{app}$ = 99 | 0.00018 | 0.0022 | 0.026 | 0.24 | 0.22 | 0.43 | 0.071 | 0.0031 |
| | 20 $K_{app}$ = 660 | 0.00026 | 0.0033 | 0.12 | 0.19 | 0.042 | 0.55 | 0.090 | 0.0039 |
| | 3 $K_{app}$ + Load | 0.000074 | 0.0009 | 0.013 | 0.12 | 0.27 | 0.53 | 0.071 | 0.0012 |
| **H251N** | | | | | | | | | |
| | $K_{app}$ = 34 | 0.00016 | 0.0040 | 0.011 | 0.26 | 0.45 | 0.22 | 0.060 | 0.0026 |
| | 3 $K_{app}$ = 102 | 0.00024 | 0.0063 | 0.035 | 0.31 | 0.22 | 0.33 | 0.091 | 0.0038 |
| | 20 $K_{app}$= 680 | 0.00035 | 0.0098 | 0.17 | 0.24 | 0.042 | 0.42 | 0.11 | 0.0049 |
| | 3 $K_{app}$ + Load | 0.00011 | 0.0028 | 0.018 | 0.16 | 0.29 | 0.43 | 0.098 | 0.0017 |
| **D382Y** | | | | | | | | | |
| | $K_{app}$ = 52 | 0.00016 | 0.0020 | 0.012 | 0.19 | 0.46 | 0.29 | 0.052 | 0.0024 |
| | 3 $K_{app}$ = 156 | 0.00025 | 0.0031 | 0.039 | 0.23 | 0.22 | 0.43 | 0.077 | 0.0036 |
| | 20 $K_{app}$ = 1040 | 0.00035 | 0.0047 | 0.19 | 0.18 | 0.039 | 0.49 | 0.089 | 0.0041 |
| | 3 $K_{app}$ + Load | 0.00010 | 0.0013 | 0.019 | 0.11 | 0.27 | 0.52 | 0.077 | 0.0015 |
| **P710R** | | | | | | | | | |
| | $K_{app}$ = 14 | 0.000062 | 0.0012 | 0.0050 | 0.27 | 0.41 | 0.28 | 0.025 | 0.0012 |
| | 3 $K_{app}$ = 42 | 0.000095 | 0.0018 | 0.015 | 0.31 | 0.20 | 0.42 | 0.038 | 0.0019 |
| | 20 $K_{app}$ = 280 | 0.00014 | 0.0029 | 0.082 | 0.29 | 0.039 | 0.54 | 0.048 | 0.0024 |
| | 3 $K_{app}$ + Load | 0.00004 | 0.00077 | 0.0080 | 0.17 | 0.25 | 0.53 | 0.038 | 0.00077 |

| | | | | | | | | | |
|---|---|---|---|---|---|---|---|---|---|
| **V763M** | | | | | | | | | |
| | $K_{app}$ = 18 | 0.00012 | 0.0015 | 0.0054 | 0.27 | 0.49 | 0.20 | 0.036 | 0.0017 |
| | 3 $K_{app}$ = 54 | 0.00019 | 0.0025 | 0.017 | 0.34 | 0.26 | 0.33 | 0.058 | 0.0028 |
| | 20 $K_{app}$ = 360 | 0.00029 | 0.0040 | 0.096 | 0.33 | 0.051 | 0.44 | 0.078 | 0.0037 |
| | 3 $K_{app}$ + Load | 0.000083 | 0.0011 | 0.0088 | 0.18 | 0.33 | 0.42 | 0.060 | 0.0012 |
| **R719W** | | | | | | | | | |
| | $K_{app}$ = 33 | 0.00012 | 0.0030 | 0.0088 | 0.21 | 0.45 | 0.30 | 0.032 | 0.0020 |
| | 3 $K_{app}$ = 99 | 0.00018 | 0.0047 | 0.027 | 0.25 | 0.22 | 0.44 | 0.048 | 0.0029 |
| | 20 $K_{app}$ = 660 | 0.00026 | 0.0074 | 0.130 | 0.19 | 0.042 | 0.56 | 0.061 | 0.0037 |
| | 3 $K_{app}$ + Load | 0.000073 | 0.0020 | 0.014 | 0.13 | 0.27 | 0.54 | 0.044 | 0.0012 |
| **R723G** | | | | | | | | | |
| | $K_{app}$ = 43 | 0.00020 | 0.0038 | 0.013 | 0.25 | 0.44 | 0.26 | 0.031 | 0.0020 |
| | 3 $K_{app}$ = 129 | 0.00031 | 0.0061 | 0.041 | 0.30 | 0.22 | 0.39 | 0.046 | 0.0030 |
| | 20 $K_{app}$ = 860 | 0.00049 | 0.010 | 0.18 | 0.21 | 0.041 | 0.49 | 0.058 | 0.0038 |
| | 3 $K_{app}$ + Load | 0.00014 | 0.0027 | 0.021 | 0.16 | 0.28 | 0.50 | 0.044 | 0.0013 |
| **G741R** | | | | | | | | | |
| | $K_{app}$ = 29 | 0.00011 | 0.0021 | 0.0077 | 0.21 | 0.45 | 0.30 | 0.037 | 0.0019 |
| | 3 $K_{app}$ = 87 | 0.00017 | 0.0032 | 0.024 | 0.24 | 0.22 | 0.45 | 0.056 | 0.0028 |
| | 20 $K_{app}$ = 580 | 0.00025 | 0.0050 | 0.12 | 0.20 | 0.042 | 0.57 | 0.071 | 0.0035 |
| | 3 $K_{app}$ + Load | 0.000069 | 0.0013 | 0.012 | 0.12 | 0.27 | 0.54 | 0.054 | 0.0011 |

**Table S5. Resolution matrices.**

| ***β-WT*** | $K_A$ | $k_{Pi}$ | $k_{-T}$ | $K_H$ | $K_{AH}$ |
|---|---|---|---|---|---|
| $K_A$ | 0.91 | 0.10 | 0.00 | -0.17 | -0.19 |
| $k_{pi}$ | 0.10 | 0.88 | 0.00 | 0.22 | 0.22 |
| $k_{-T}$ | 0.00 | 0.00 | 0.00 | 0.00 | 0.00 |
| $K_H$ | -0.17 | 0.22 | 0.00 | 0.59 | -0.39 |
| $K_{AH}$ | -0.19 | 0.22 | 0.00 | -0.39 | 0.59 |

| **H251N** | $K_A$ | $k_{Pi}$ | $k_{-T}$ | $K_H$ | $K_{AH}$ |
|---|---|---|---|---|---|
| $K_A$ | 0.79 | 0.22 | 0.00 | -0.23 | -0.24 |
| $k_{pi}$ | 0.22 | 0.75 | 0.00 | 0.26 | 0.26 |
| $k_{-T}$ | 0.00 | 0.00 | 0.00 | 0.00 | 0.00 |
| $K_H$ | -0.23 | 0.26 | 0.00 | 0.72 | -0.27 |
| $K_{AH}$ | -0.24 | 0.26 | 0.00 | -0.27 | 0.72 |

| **D382Y** | $K_A$ | $k_{Pi}$ | $k_{-T}$ | $K_H$ | $K_{AH}$ |
|---|---|---|---|---|---|
| $K_A$ | 0.91 | 0.09 | 0.00 | -0.17 | -0.18 |
| $k_{pi}$ | 0.09 | 0.89 | 0.00 | 0.21 | 0.21 |
| $k_{-T}$ | 0.00 | 0.00 | 0.00 | 0.00 | 0.00 |
| $K_H$ | -0.17 | 0.21 | 0.00 | 0.58 | -0.40 |
| $K_{AH}$ | -0.18 | 0.21 | 0.00 | -0.40 | 0.58 |

| P710R | $K_A$ | $k_{Pi}$ | $k_{-T}$ | $K_H$ | $K_{AH}$ |
|---|---|---|---|---|---|
| $K_A$ | 0.88 | 0.12 | 0.00 | -0.17 | -0.19 |
| $k_{pi}$ | 0.12 | 0.84 | 0.00 | 0.24 | 0.25 |
| $k_{-T}$ | 0.00 | 0.00 | 0.00 | 0.00 | 0.00 |
| $K_H$ | -0.17 | 0.24 | 0.00 | 0.62 | -0.36 |
| $K_{AH}$ | -0.19 | 0.25 | 0.00 | -0.36 | 0.62 |

| V763M | $K_A$ | $k_{Pi}$ | $k_{-T}$ | $K_H$ | $K_{AH}$ |
|---|---|---|---|---|---|
| $K_A$ | 0.81 | 0.21 | 0.00 | -0.22 | -0.23 |
| $k_{pi}$ | 0.21 | 0.75 | 0.00 | 0.26 | 0.27 |
| $k_{-T}$ | 0.00 | 0.00 | 0.00 | 0.00 | 0.00 |
| $K_H$ | -0.22 | 0.26 | 0.00 | 0.72 | -0.28 |
| $K_{AH}$ | -0.23 | 0.27 | 0.00 | -0.28 | 0.71 |

| R719W | $K_A$ | $k_{Pi}$ | $k_{-T}$ | $K_H$ | $K_{AH}$ |
|---|---|---|---|---|---|
| $K_A$ | 0.90 | 0.10 | 0.00 | -0.17 | -0.19 |
| $k_{pi}$ | 0.10 | 0.87 | 0.00 | 0.22 | 0.22 |
| $k_{-T}$ | 0.00 | 0.00 | 0.00 | 0.00 | 0.00 |
| $K_H$ | -0.17 | 0.22 | 0.00 | 0.60 | -0.39 |
| $K_{AH}$ | -0.19 | 0.22 | 0.00 | -0.39 | 0.59 |

| R723G | $K_A$ | $k_{pi}$ | $k_{-T}$ | $K_H$ | $K_{AH}$ |
|---|---|---|---|---|---|
| $K_A$ | 0.60 | 0.42 | 0.00 | -0.16 | -0.17 |
| $k_{pi}$ | 0.42 | 0.56 | 0.00 | 0.18 | 0.18 |
| $k_{-T}$ | 0.00 | 0.00 | 0.00 | 0.00 | 0.00 |
| $K_H$ | -0.16 | 0.18 | 0.00 | 0.93 | -0.07 |
| $K_{AH}$ | -0.17 | 0.18 | 0.00 | -0.07 | 0.92 |

| G741R | $K_A$ | $k_{pi}$ | $k_{-T}$ | $K_H$ | $K_{AH}$ |
|---|---|---|---|---|---|
| $K_A$ | 0.91 | 0.09 | 0.00 | -0.16 | -0.18 |
| $k_{pi}$ | 0.09 | 0.89 | 0.00 | 0.21 | 0.22 |
| $k_{-T}$ | 0.00 | 0.00 | 0.00 | 0.00 | 0.00 |
| $K_H$ | -0.16 | 0.21 | 0.00 | 0.59 | -0.40 |
| $K_{AH}$ | -0.18 | 0.22 | 0.00 | -0.40 | 0.58 |

**Table S6. Sensitivity analysis; Percentage change of the six fitted parameters induced by a change of +20% or -20% to one fitted parameters for the P710R and R723G mutations.**

| ***Equilibrium Rate Constants*** | ***Units*** | ***ß-R723G*** | **$V_{max}$+20%** | **$V_{max}$ -20%** | **$K_{D*}$ +20%** | **$K_{D*}$ -20%** | **$K_H$ +20%** | **$K_H$ -20%** |
|---|---|---|---|---|---|---|---|---|
| $K_m$ | (μM) | 43 | 0.0 | 0.0 | 0.0 | 0.0 | 0.0 | 0.0 |
| $V_{max}$ | $s^{-1}$ | 4 | 20.0 | -20.0 | 0.0 | 0.0 | 0.0 | 0.0 |
| $K_A$ | $\mu M^{-1}$ | 0.014 | 1.4 | 14.3 | 2.1 | -2.1 | 10.7 | -16.4 |
| $K_{pi}$ | mM | 100 | 0.0 | 0.0 | 0.0 | 0.0 | 0.0 | 0.0 |
| $K_{D*}$ | | 0.167 | 0.0 | 0.0 | 20.0 | -20.0 | 0.0 | 0.0 |
| $K_D$ | (μM) | 36 | 0.0 | 0.0 | 0.0 | 0.0 | 0.0 | 0.0 |
| $K_T$ | $\mu M^{-1}$ | 0.0049 | 0.0 | 0.0 | 0.0 | 0.0 | 0.0 | 0.0 |
| $K_{T*}$ | | 55.6 | 0.0 | 0.0 | 0.0 | 0.0 | 0.0 | 0.0 |
| $K_{T**}$ | μM | 1000 | 0.0 | 0.0 | 0.0 | 0.0 | 0.0 | 0.0 |
| $K_H$ | | 7.2 | 30.5 | -7.2 | -0.2 | 0.3 | 20.0 | -20.0 |
| $K_{AH}$ | | 51.4 | 30.6 | -7.1 | -0.2 | 0.3 | 20.2 | -20.1 |
| ***Forward Rate Constants*** | | | | | | | | |
| $k_A$ | $\mu M^{-1}s^{-1}$ | 14 | 1.4 | 14.3 | 2.1 | -2.1 | 10.7 | -16.4 |
| $k_{Pi}$ | $s^{-1}$ | 7.7 | 15.6 | -29.9 | -1.3 | 2.6 | -11.7 | 23.4 |
| $k_{D*}$ | $s^{-1}$ | 106.8 | 0.0 | 0.0 | 20.0 | -20.0 | 0.0 | 0.0 |
| $k_D$ | $s^{-1}$ | 1000 | 0.0 | 0.0 | 0.0 | 0.0 | 0.0 | 0.0 |
| $k_T$ | $\mu M^{-1}s^{-1}$ | 10.07 | 0.0 | 0.0 | 0.0 | 0.0 | 0.2 | -0.1 |
| $k_{T*}$ | $s^{-1}$ | 556 | 0.0 | 0.0 | 0.0 | 0.0 | 0.0 | 0.0 |
| $k_{T**}$ | $s^{-1}$ | 1000 | 0.0 | 0.0 | 0.0 | 0.0 | 0.0 | 0.0 |
| $k_H$ | $s^{-1}$ | 10.0 | 30.5 | -7.2 | -0.2 | 0.3 | 20.0 | -20.0 |
| $k_{AH}$ | $s^{-1}$ | 10.3 | 30.6 | -7.1 | -0.2 | 0.3 | 20.2 | -20.1 |

| Backward Rate Constants | | | | | | | | |
|---|---|---|---|---|---|---|---|---|
| $k_{-A}$ | $s^{-1}$ | 1000 | 0.0 | 0.0 | 0.0 | 0.0 | 0.0 | 0.0 |
| $k_{-Pi}$ | $mM^{-1}s^{-1}$ | 0.077 | 15.6 | -29.9 | -1.3 | 2.6 | -11.7 | 23.4 |
| $k_{-D*}$ | $s^{-1}$ | 639.52 | 0.0 | 0.0 | 0.0 | 0.0 | 0.0 | 0.0 |
| $k_{-D}$ | $\mu M^{-1}s^{-1}$ | 27.78 | 0.0 | 0.0 | 0.0 | 0.0 | 0.0 | 0.0 |
| $k_{-T}$ | $s^{-1}$ | 2054.3 | 0.0 | 0.0 | 0.0 | 0.0 | 0.2 | -0.1 |
| $k_{-T*}$ | $s^{-1}$ | 10 | 0.0 | 0.0 | 0.0 | 0.0 | 0.0 | 0.0 |
| $k_{-T**}$ | $\mu M^{-1}s^{-1}$ | 1 | 0.0 | 0.0 | 0.0 | 0.0 | 0.0 | 0.0 |
| $k_{-H}$ | $s^{-1}$ | 1.4 | 0.0 | 0.0 | 0.0 | 0.0 | 0.0 | 0.0 |
| $k_{-AH}$ | $s^{-1}$ | 0.2 | 0.0 | 0.0 | 0.0 | 0.0 | 0.0 | 0.0 |

B.

| ***Equilibrium Rate Constants*** | ***Units*** | ***P710R*** | $V_{max}$+20% | $V_{max}$ -20% | $K_{D*}$ +20% | $K_{D*}$ -20% | $K_H$ +20% | $K_H$ -20% |
|---|---|---|---|---|---|---|---|---|
| $K_m$ | (μM) | 14 | 0.0 | 0.0 | 0.0 | 0.0 | 0.0 | 0.0 |
| $V_{max}$ | $s^{-1}$ | 2.5 | 20.0 | -20.0 | 0.0 | 0.0 | 0.0 | 0.0 |
| $K_A$ | $\mu M^{-1}$ | 0.0501 | -7.0 | 6.6 | 1.4 | -0.6 | 8.2 | -13.2 |
| $K_{pi}$ | mM | 100 | 0.0 | 0.0 | 0.0 | 0.0 | 0.0 | 0.0 |
| $K_{D*}$ | | 0.167 | 0.0 | 0.0 | 20.0 | -20.0 | 0.0 | 0.0 |
| $K_D$ | (μM) | 36 | 0.0 | 0.0 | 0.0 | 0.0 | 0.0 | 0.0 |
| $K_T$ | $\mu M^{-1}$ | 0.0062 | 0.0 | 0.0 | 0.0 | 0.0 | 0.0 | 0.0 |
| $K_{T*}$ | | 110.65 | 0.0 | 0.0 | 0.0 | 0.0 | 0.0 | 0.0 |
| $K_{T**}$ | μM | 1000 | 0.0 | 0.0 | 0.0 | 0.0 | 0.0 | 0.0 |
| $K_H$ | | 4.9 | 13.6 | -18.2 | -0.2 | 4.0 | 20.0 | -20.0 |
| $K_{AH}$ | | 34.7 | 13.6 | -18.2 | -0.2 | 4.1 | 20.4 | -20.3 |

| | | | | | | | | |
|---|---|---|---|---|---|---|---|---|
| ***Forward Rate Constants*** | | | | | | | | |
| $k_A$ | $\mu M^{-1}s^{-1}$ | 50.1 | -7.0 | 6.6 | 1.4 | -0.6 | 8.2 | -13.2 |
| $k_{Pi}$ | $s^{-1}$ | 4.4 | 27.3 | -20.5 | 0.0 | 1.1 | -9.1 | 20.5 |
| $k_{D*}$ | $s^{-1}$ | 69 | 0.0 | 0.0 | 20.0 | -20.0 | 0.0 | 0.0 |
| $k_D$ | $s^{-1}$ | 1000 | 0.0 | 0.0 | 0.0 | 0.0 | 0.0 | 0.0 |
| $k_T$ | $\mu M^{-1}s^{-1}$ | 10.25 | 0.0 | 0.0 | 0.1 | -0.3 | 0.1 | -1.8 |
| $k_{T*}$ | $s^{-1}$ | 1106.5 | 0.0 | 0.0 | 0.0 | 0.0 | 0.0 | 0.0 |
| $k_{T**}$ | $s^{-1}$ | 1000 | 0.0 | 0.0 | 0.0 | 0.0 | 0.0 | 0.0 |
| $k_H$ | $s^{-1}$ | 6.8 | 13.6 | -18.2 | -0.2 | 4.0 | 20.0 | -20.0 |
| $k_{AH}$ | $s^{-1}$ | 6.9 | 13.6 | -18.2 | -0.2 | 4.1 | 20.4 | -20.3 |
| ***Backward Rate Constants*** | | | | | | | | |
| $k_{-A}$ | $s^{-1}$ | 1000 | 0.0 | 0.0 | 0.0 | 0.0 | 0.0 | 0.0 |
| $k_{-Pi}$ | $mM^{-1}s^{-1}$ | 0.044 | 27.3 | -20.5 | 0.0 | 1.1 | -9.1 | 20.5 |
| $k_{-D*}$ | $s^{-1}$ | 413.17 | 0.0 | 0.0 | 0.0 | 0.0 | 0.0 | 0.0 |
| $k_{-D}$ | $\mu M^{-1}s^{-1}$ | 27.78 | 0.0 | 0.0 | 0.0 | 0.0 | 0.0 | 0.0 |
| $k_{-T}$ | $s^{-1}$ | 1653.5 | 0.0 | 0.0 | 0.1 | -0.3 | 0.1 | -1.8 |
| $k_{-T*}$ | $s^{-1}$ | 10 | 0.0 | 0.0 | 0.0 | 0.0 | 0.0 | 0.0 |
| $k_{-T**}$ | $\mu M^{-1}s^{-1}$ | 1 | 0.0 | 0.0 | 0.0 | 0.0 | 0.0 | 0.0 |
| $k_{-H}$ | $s^{-1}$ | 1.4 | 0.0 | 0.0 | 0.0 | 0.0 | 0.0 | 0.0 |
| $k_{-AH}$ | $s^{-1}$ | 0.2 | 0.0 | 0.0 | 0.0 | 0.0 | 0.0 | 0.0 |

**Table S7. Balance of significant rate constants around the ATPase cycle.**

| Isoform | $k_{cat}$ ($s^{-1}$) | $k_{Pi}/k_{cat}$ | $k_{D*}/k_{cat}$ | $k_H/k_{cat}$ |
|---|---|---|---|---|
| WT | 4.1 | 1.7 | 14.4 | 3.2 |
| H251N | 5.1 | 2.3 | 11.1 | 2.4 |
| D382Y | 4.9 | 1.7 | 13.1 | 3.3 |
| P710R | 2.5 | 1.8 | 27.6 | 2.7 |
| V763M | 4 | 2.2 | 17 | 2.3 |
| R719W | 3.9 | 1.7 | 24.8 | 3.1 |
| R723G | 4 | 1.9 | 26.7 | 2.5 |
| G741R | 3.7 | 1.7 | 19.2 | 3.2 |

**Table S8. Comparison of velocity data for wild-type and HCM mutant β-cardiac sS1 at 23 °C.**

| Property | **WT** | H251N[#] | D382Y** | P710R** | V763M** | **WT** | R719W* | **WT** | R723G* | **WT** | G741R* |
|---|---|---|---|---|---|---|---|---|---|---|---|
| MVIS (nm/s) | 858 ± 11 | 1195 ± 16 | 769 ± 14 | 275 ± 15 | | 596 ± 21 | 606 ± 22 | 646 ± 25 | 686 ± 26 | 579 ± 48 | 589 ± 49 |
| TOP5% (nm/s) | 1213 ± 15 | 1729 ± 34 | 1143 ± 23 | 468 ± 27 | 1363 ± 24 | 893 ± 26 | 1019 ± 28 | 958 ± 32 | 1068 ± 37 | 905 ± 63 | 885 ± 65 |
| Rel. MVIS | 1 | 1.3927 | 0.8962 | 0.3205 | | 1 | 1.0167 | 1 | 1.0619 | 0.8962 | 0.9117 |
| Rel. Top 5% | 1 | 1.4254 | 0.9423 | 0.3858 | 1.1236 | 1 | 1.1410 | 1 | 1.1148 | 0.9446 | 0.9238 |

Mean ± SEM from the number of motor preparations.
MVIS is the mean velocity including stuck filaments.
TOP 5% is the mean of top 5% of all the velocity data.
Relative values are also tabulated to highlight the difference.

= means those mutations were compared side-by-side to WT proteins that were cultured simultaneously and tested on the same flow chamber.

[#] = Published in (3)    * = Published in (4)    ** = Unpublished data

Filaments that are tracked longer than 10 frames are included in the analysis. A 20% tolerance filter is applied to eliminate intermittently moving filaments with a velocity dispersion higher than 20% of their mean within a 5-frame window. Each velocity point in the analysis is the average velocity over 5 frames. At least four replicate movies from the indicated number of different preparations of sS1 were analysed.

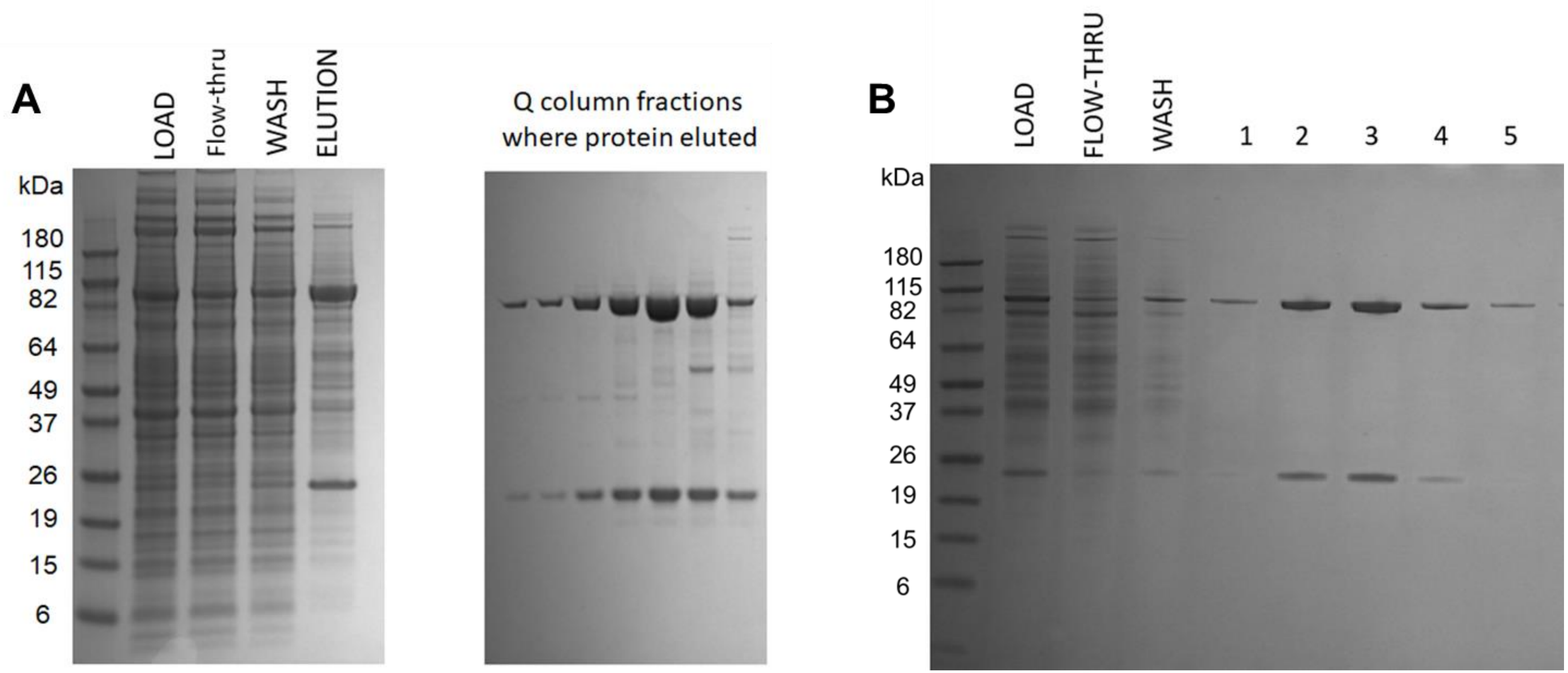

**Figure S1.** (A) Representative gels from a purification of β-R719W. Left is a His-NTA column by gravity flow starting with the lysate and ending with the imidazole eluted sS1. On the right is part of a gel with the sS1 containing fractions. (B) is a representative gel from a purification of β-V763M. After a gravity His column, the solution was loaded on a PDZ column, where the collected fractions also resulted in clean sS1 (sS1: 93 kDa, MYL3: 25 kDa).

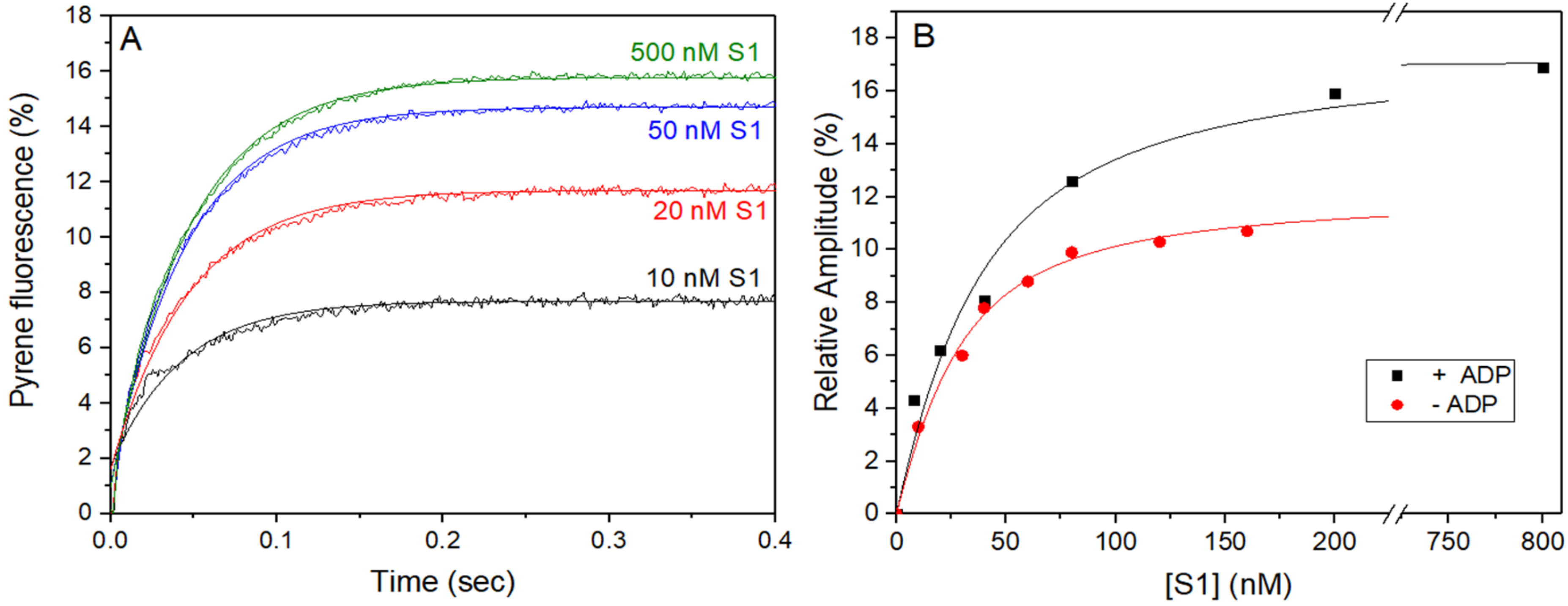

**Figure S2. H251N sS1 affinity for actin in the presence and absence of ADP.** (A) Example traces of the increase in pyrene fluorescence when 10 µM ATP is mixed with 15 nM pyr-actin, 200 µM ADP and varying concentrations of sS1 (10-500 nM). The pyr-actin was treated with hexokinase and $Ap_5A$ to ensure no ATP was present in the protein before mixing in the stopped-flow. (B) Fluorescence amplitude plotted as a function of sS1 concentration can be described by a quadratic fit resulting in a $K_{DA}$ of 65.6 ± 13 nM in the presence of ADP, and a $K_A$ of 11.5 ± 1.8 nM when in the absence of ADP. The average values from 3 independent measurements are given in Table 1.

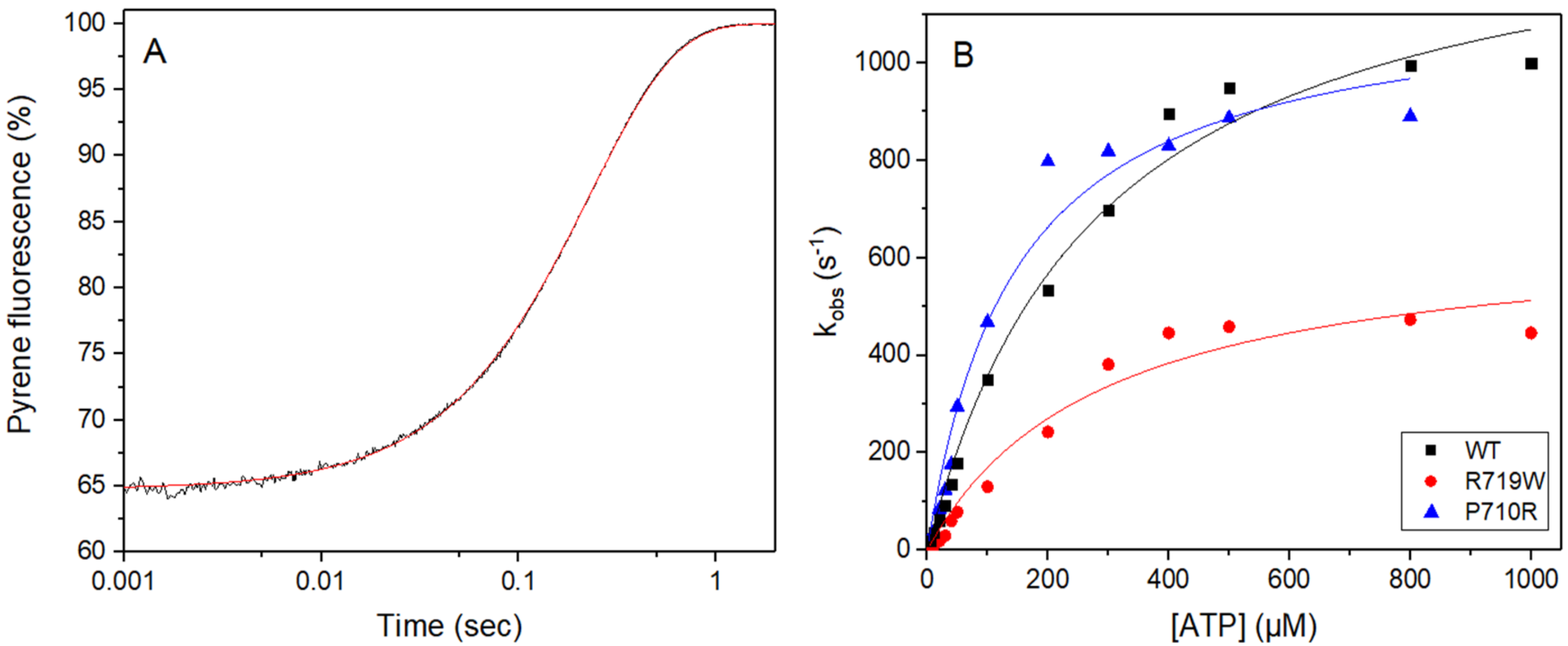

**Figure S3. ATP induced-dissociation of acto.sS1.** (A) Example stopped-flow trace to measure ATP-induced dissociation of pyr-actin from the WT sS1. 10 μM ATP was mixed with 50 nM pyr-actin.sS1, and the increase in pyrene fluorescence was measured. The experiment was repeated for all constructs and the observed amplitude of the fluorescence changes were similar (within 30%) in each case. (B) The effect of ATP concentration on k for ATP-induced dissociation of pyr-actin.sS1 for 3 proteins; WT, R719W and P710R sS1. The gradient of the initial slope generates a second order rate constant of ATP binding, $K_T k_{+T^*}$. The best fit to the hyperbola also yields a maximum k = $k_{+T^*}$ and the ATP concentration required for half maximum k = $K_T$. Best fit values for 3 independent measurements of all constructs are in Table 1.

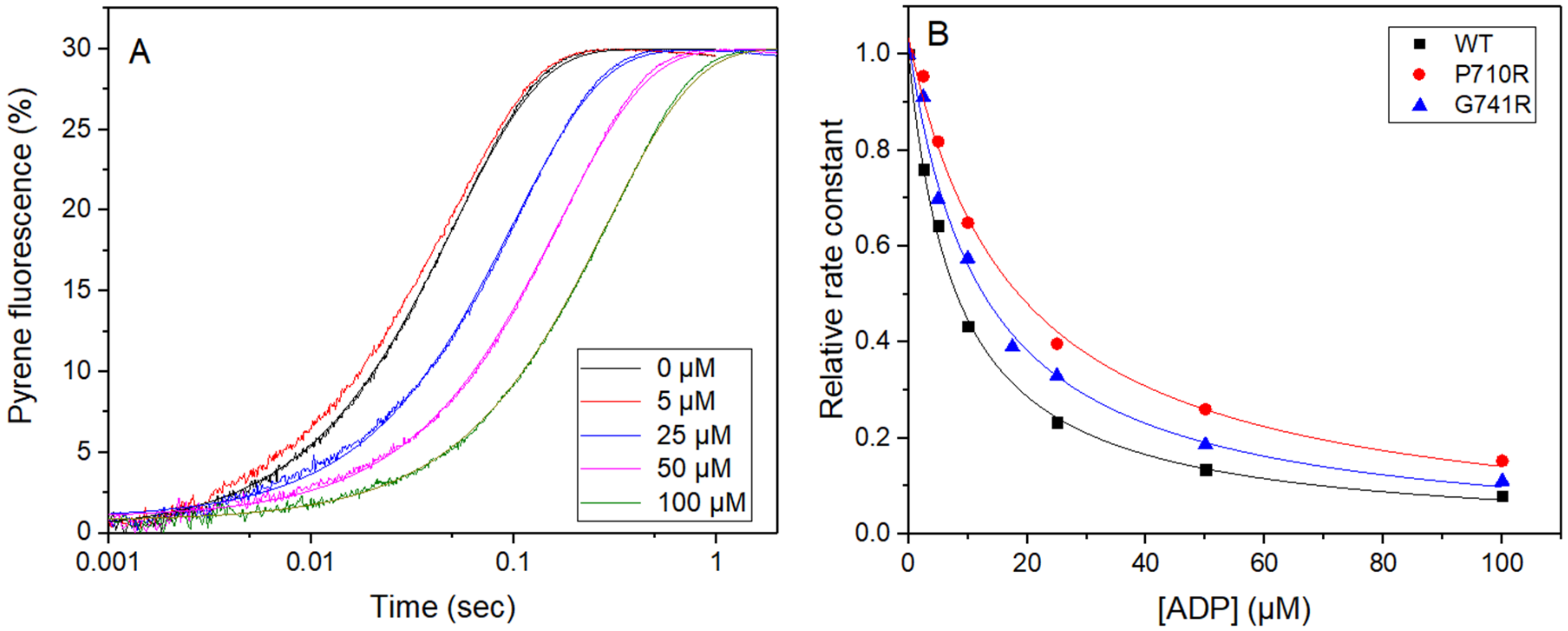

**Figure S4. ADP affinity for acto.sS1.** (A) Example transients showing an increase in pyrene fluorescence when 100 nM pyr-actin.WT sS1 is mixed with 10 µM ATP and 0-100 mM ADP. (B) k plotted as a function of ADP concentration for WT, P710R and G741R sS1 proteins, showing hyperbolic dependence. The value of $((K_{D^*}+1)/K_DK_{D^*})$ is given by the ADP concentration at 50% inhibition.

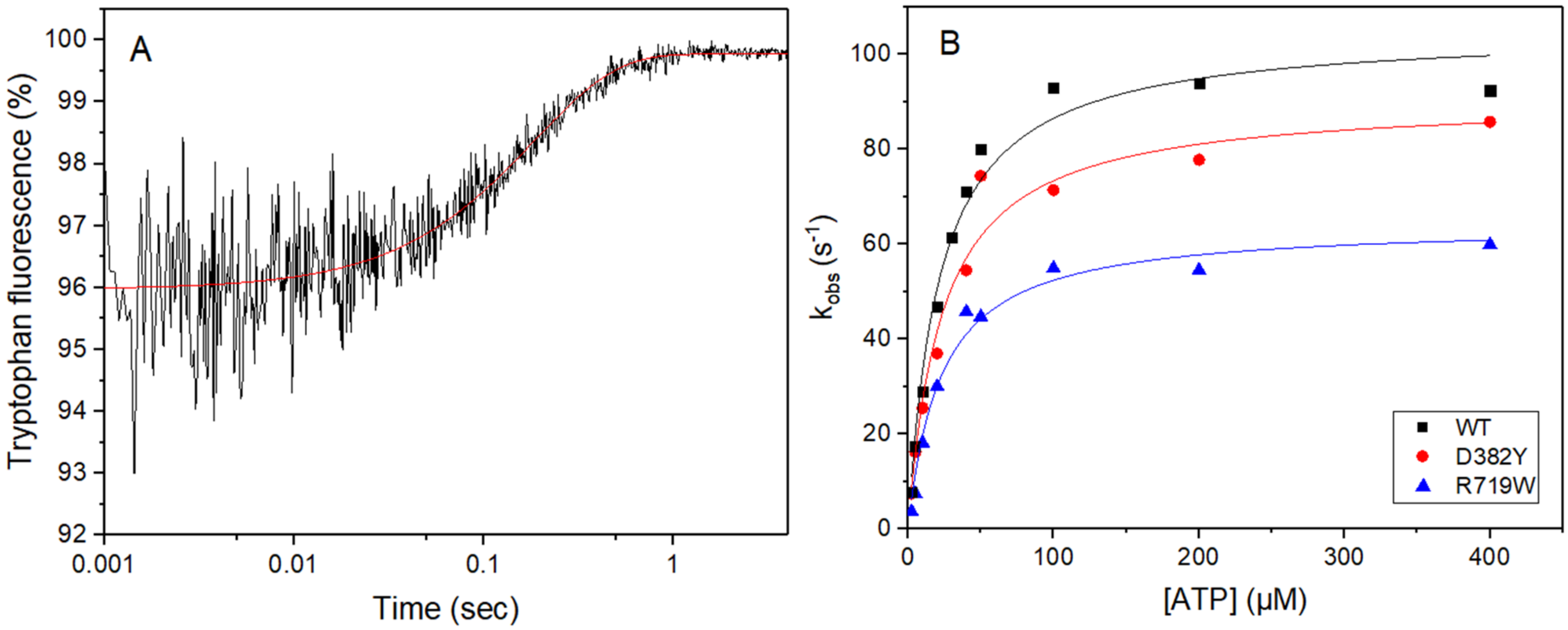

**Figure S5. ATP binding to sS1.** (A) Example stopped-flow trace to measure ATP binding to WT sS1. 10 μM ATP was mixed with 200 nM sS1, and the increase in tryptophan fluorescence was measured. The sS1 was pre-treated with apyrase to ensure no nucleotide was present before mixing in the stopped-flow. Least squares best fit to a single exponential is shown in red with k = 22 $s^{-1}$ and a 4% increase in pyrene fluorescence. (B) The hyperbolic dependence of ATP concentration on k. The best fit to a hyperbola yields max k = $k_{+H}$ + $k_{--H}$, and the ATP concentration required for half maximum k = $K_{50\%}$. The initial slope defines the second order rate constant of ATP binding to sS1. Best fit values for 3 independent measurements of all constructs are in Table 1.

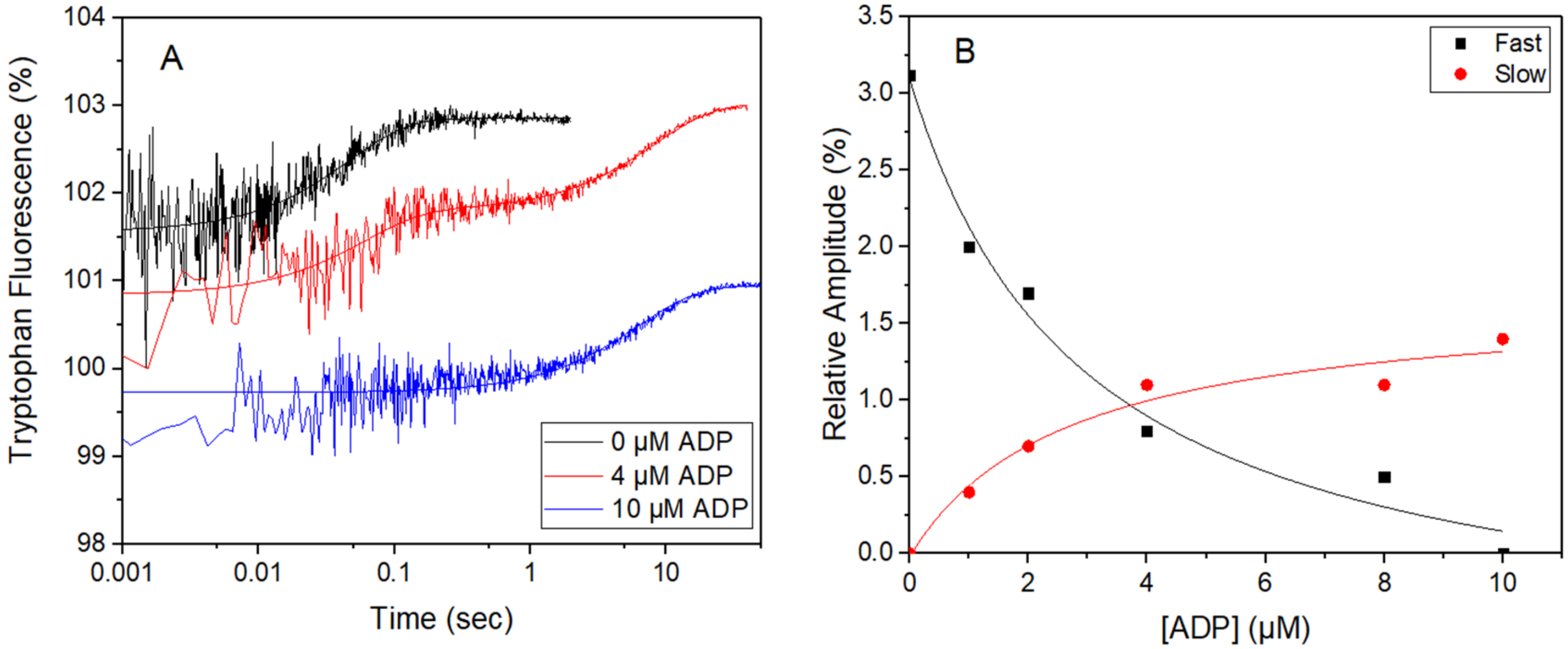

**Figure S6. ADP affinity for P710R sS1**. (A) Example traces of 50 μM ATP mixed with 200 nM sS1 that is preincubated with ADP. The black trace contains no ADP, and results in a transient best described by a single exponential fit. In the presence of 4 μM ADP, the amplitude of the fast phase decreased as the concentration of free sS1 decreases, and a second slow phase is observed representing the release of ADP from sS1. The transient is best described by a double exponential. At high ADP concentration, the amplitude of the fast phase is lost, and the amplitude of the slow phase can be best described by a single exponential. (B) The relative amplitudes of the fast and slow phases plotted against ADP concentration, which has a hyperbolic dependence. Table 1 lists averages for 3 independent measurements for all constructs.
